# Speciation completion rates have limited impact on macroevolutionary diversification

**DOI:** 10.1101/2024.07.01.601000

**Authors:** Pierre Veron, Jérémy Andréoletti, Tatiana Giraud, Hélène Morlon

**Affiliations:** Institut de Biologie, École Normale Supérieure, Université PSL, CNRS, INSERM, Paris, France; Écologie Systématique et Évolution, CNRS, Université Paris-Saclay, AgroParisTech, Gif-sur-Yvette, France

**Keywords:** speciation, macroevolution, microevolution, phylogeny

## Abstract

Standard birth-death processes used in macroevolutionary studies assume instantaneous speciation, an unrealistic premise that limits the interpretation of speciation and extinction rates. The protracted birth-death (PBD) model instead assumes that speciation involves two steps: initiation and completion. In order to understand their respective influence on macroevolutionary speciation rates, we compute a standard time-varying birth-death scenario that is “equivalent” to the PBD model in terms of speciation and extinction probabilities. First, we find a sharp decline in the equivalent birth rate near the present, indicating that rates estimated at the tips of phylogenies may not accurately reflect the underlying speciation process. Second, the completion rate controls the timing of the decay rather than the asymptotic equivalent rates. The equivalent birth rate in the past scales with the speciation initiation rate, with a scaling factor depending mostly on the population extinction rate. Our results suggest that the rates of population formation and extinction may often play a larger role than the speed of accumulation of reproductive isolation in modulating speciation rates. Our study establishes a theoretical framework for understanding how microevolutionary processes combine to explain the diversification of species on macroevolutionary time scales.

## 1 Introduction

Birth-death models are widely used to understand the diversification of species groups; in this context, births represent speciation events, i.e. the emergence of two daughter species from an ancestral one, and deaths represent species extinction. This specific use of birth-death models is particularly widespread to interpret both fossil (Silvestro et al. 2014) and phylogenetic data (Stadler 2013; Morlon et al. 2024) in terms of diversification dynamics. In birth-death models, speciation is considered to be an instantaneous phenomenon, represented as a branching event following a Poisson point process.

Despite the widespread use of birth-death models to represent speciation and extinction events, speciation is not instantaneous. Speciation requires an initial isolation of populations (speciation initiation) followed by the accumulation of genetic barriers to gene flow until speciation is complete. Speciation can initiate and fail before completion, for example because of secondary contact, or because isolated populations go extinct before speciation completion (Coyne and Orr 2004; Rosenblum et al. 2012; Dynesius and Jansson 2014). The whole speciation process may take hundreds of thousands up to several millions of years (Benton and Pearson 2001; Etienne and Rosindell 2012; Etienne et al. 2014; Hua et al. 2022).

Ignoring the fact that speciation takes time by using standard birth-death models has non trivial consequences for our understanding of diversification dynamics. For example, when standard birth-death models are used in combination with phylogenetic trees of extant species to estimate speciation and extinction rates, the “protracted” nature of speciation may be misinterpreted as a speciation rate slowdown towards the present (Etienne and Rosindell 2012; Moen and Morlon 2014).

Etienne and Rosindell 2012 pioneered the development of the so-called protracted birth-death model (PBD). Instead of assuming that speciation is instantaneous as in the standard birth-death model, this model assumes that there are events of speciation *initiation* corresponding to the formation of *incipient* species that eventually become *good* species after a random *completion* time. An incipient lineage is a lineage that is not yet considered as a different species from the ancestral lineage. For sexually reproducing organisms, the completion time is the time it takes for lineages to achieve reproductive isolation. Each lineage is thus subject to initiation, extinction and completion events. Although the PBD model is phenomenological, the protracted nature of speciation represents the effect of mechanisms such as the accumulation of genetic incompatibilities between lineages (for instance, Bateson-Dobzhanski-Muller incompatibilities), gene flow between incipient species and adaptation that are not explicit in this model but contribute to modulate the completion time. (Etienne et al. 2014).

The protracted birth-death model has several advantages over the the standard birthdeath model. First, it is biologically more realistic, and it thus unsurprisingly produces phylogenies that are closer to empirical phylogenies than those produced by the standard birth-death model (Etienne and Rosindell 2012). Specifically, it produces phylogenies that are less tippy (fewer recent speciation events) than those arising from the standard birth-death model. Second, it allows the integration of intraspecific processes that lead to speciation. For example, the matching competition birth-death model (MCBD, Aristide and Morlon 2019) integrates the effect of intraspecific competition on character displacement leading to speciation by modeling character displacement in incipient lineages.

Despite the advantages of the protracted birth-death model, the overwhelming majority of phylogenetic analyses of diversification use the standard birth-death model. Most available models for phylogenetic analyses of diversification are versions of the standard birth-death model, with birth and death rates that can vary in time and/or across lineages (Morlon et al. 2024). The protracted birth-death model can be fitted to empirical phylogenies; however not all of its parameters can be reliably estimated from a phylogeny (Etienne et al. 2014), which limits its usefulness. Recently, Hua et al. showed that the parameters of a protracted speciation model (slightly different from PBD) can be accurately estimated from population-level (rather than species-level) phylogenies; however such phylogenies remain rare (Hua et al. 2022). Fitting standard birth-death models thus remains the norm in phylogenetic analyses of diversification.

If speciation takes time but is estimated by fitting standard birth-death models to phylogenies, which assume instantaneous speciation, what do resulting speciation and extinction rate estimates actually represent? We can expect that speciation rate estimates will be higher when rates of speciation initiation and completion are higher, and rates of extinction of incipient species are lower, but precisely answering this question requires to establish an analytical relationship between the parameters of the protracted and standard birth-death models. To our knowledge, such a relationship has not yet been established.

Elucidating the relationship between the parameters of the protracted and standard birth-death models is important not only to clarify the meaning of speciation rates estimated from phylogenies, but also to understand the microevolutionary processes that modulate these rates (Morlon et al. 2024). Indeed, macroevolutionary speciation rates (estimated from phylogenies) vary by orders of magnitude (Maliet et al. 2019; Quintero et al. 2024), but the processes underlying this variation remain unclear. Efforts to find empirical correlations between macroevolutionary speciation rates and rates of population formation or evolution of reproductive isolation have not been conclusive; a proposed explanation is that this expected correlation is erased by the frequent extinction of incipient species (Rabosky and Matute 2013; Singhal et al. 2022; Singhal et al. 2018). The idea is that, population survival rather than population formation and the accumulation of reproductive barriers may be the factor “limiting” speciation. More generally, each of speciation initiation, speciation completion and population survival may be the process limiting macroevolutionary speciation rates in some situations but not others (Rabosky 2016). For instance, a lineage that has a propensity to accumulate fast reproductive isolation but does not experience frequent population splits might not have a high speciation rate: here, the rate of speciation completion is not limiting. Hence, factors acting on speciation initiation, the accumulation of reproductive isolation and the extinction of incipient species can have non trivial outcomes in terms of speciation rate.

Here, we obtain a mathematical link between the parameters of the protracted and standard birth-death diversification models by computing “equivalent” speciation and extinction rates, meant to represent the macroevolutionary outcomes (macroevolutionary speciation and extinction rates) of the protracted birth-death process. More precisely, our equivalent rates generate the same speciation and extinction probabilities as the protracted process. As we discuss in the paper, this is distinct from computing “congruent” (sensu Louca and Pennell 2020) speciation and extinction rates that would provide the same likelihood of extant species trees. Our main interest here is on the speciation and extinction rate outcomes of the protracted birth-death process, beyond what can be inferred from phylogenies.

## 2 Materials and methods

### 2.1 Birth-death and protracted birth-death models

The standard birth-death (BD) model involves two rates (figure 1**A**): the birth (speciation) rate *λ* and the death (extinction) rate *µ*. The protracted birth-death (PBD) model as defined in (Etienne et al. 2014) involves 5 rates (figure 1**B**) : the rate of speciation initiation from a good species *λ*_1_, the rate of completion *λ*_2_, the rate of speciation initiation from an incipient species *λ*_3_, the rate of extinction of a good species *µ*_1_ and the rate of extinction of an incipient species *µ*_2_.

**Figure 1.**
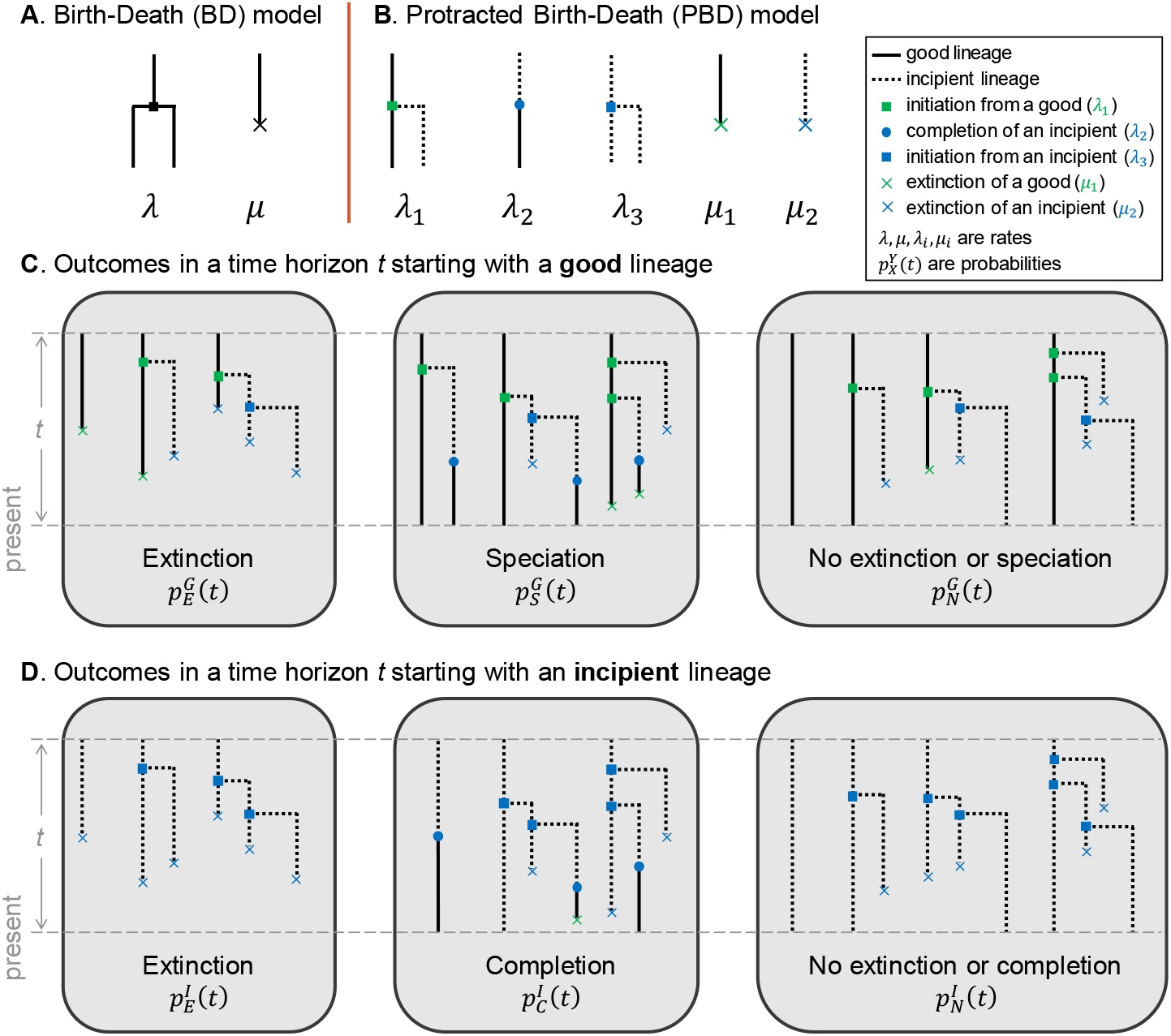
The birth-death (BD) and protracted birth-death (PBD) models. (**A, B**) Illustrations of the rates involved in the BD model (**A**), and the PBD model (**B**). (**C, D**) Possible outcomes of the PBD process in a fixed time horizon, starting from a good lineage (**C**) or an incipient lineage (**D**); non exhaustive examples of events leading to these outcomes are shown.

By convention, time elapses in the direction of increasing values. The process begins at time 0 with one good lineage and runs until the present at time *T*. We now consider an intermediate time *T* − *t*, and introduce 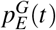 and 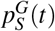 the probabilities that, for any given good lineage, the first event is an extinction or a speciation event, respectively, and 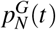 the probability that none of these events occur during a time interval of length *t* (figure 1**C**). Similarly, 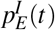 and 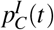 are the probabilities that, during a time interval of length *t*, the first event is respectively an extinction, or a completion event, when starting with an incipient lineage; 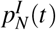 is the probability that none of these events occur during the time interval (figure 1**D**). By “extinction” we mean the direct extinction of the lineage in question, or the extinction of all possible descendants of this lineage in the time interval considered. Similarly, by “completion” we mean the direct completion of the lineage in question, or the completion of an incipient daughter lineage in the time interval under consideration. In case of multiple events occurring in the time interval (e.g. speciation followed by extinction), we consider only the first event. For instance the third case of speciation in figure 1**C** (where the lineages all go extinct after speciation) is considered as a speciation event and is recorded in 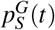. We only consider the events of speciation and extinction and not the intermediate steps of speciation initiation, because we are interested in comparing these outcomes with those of the standard birth-death process. In particular, an initiation event followed by an extinction of the two lineages will be considered as an extinction. We will however keep track of the initiation events when computing the speciation and extinction probabilities.

### 2.2 Equivalent time-dependent BD rates

#### Definitions of equivalent birth and death rates, and relation with probabilities of speciation and extinction under the PBD

Given a PBD process with fixed parameter values running from 0 to *T*, we assume that the probabilities of speciation and extinction within an infinitesimal time interval [*t* − d*t, t*] can be written as 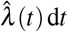 and 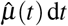, for any time *t*. We call the quantities 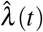 and 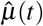 the time-dependent equivalent birth and death rates. Given a good lineage alive at time *T* − *t* − d*t*, the probability that the first event occurring within the time interval [*T* −*t* − d*t, T*] is a speciation event is given by:

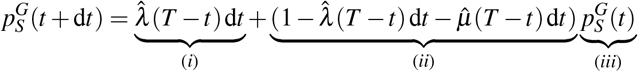

with (*i*) the probability of speciation within the small time interval [*T* −*t* − d*t, T* −*t*], (*ii*) the probability of no speciation nor extinction within the small time interval [*T* −*t* − d*t, T* −*t*] and (*iii*) the probability that the first event occurring within the time interval [*T* −*t, T*] is a speciation event, conditioned on the existence of the lineage at the time *T* −*t*. Similarly we have:

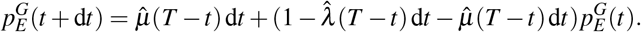

Hence, we have the dynamical system

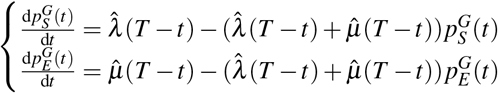

which allows us to express the time-dependent birth rate:

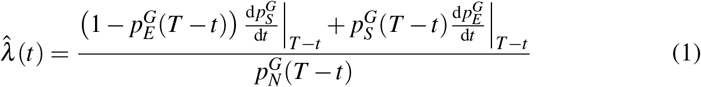

and death rate:

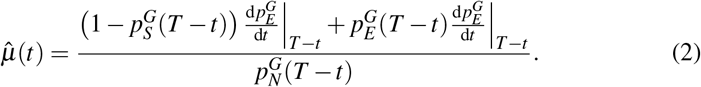

In what follows, we compute the probabilities 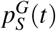 and 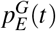 for any time *t* ∈ [0, *T*], which provides us with the equivalent rates.

By assuming that the probabilities of speciation and extinction within an infinitesimal time interval [*t* − d*t, t*] can be written as 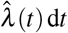 and 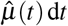, we have assumed that these probabilities depend only on time, and not on the history of the lineage considered. This is an approximation, because (under the PBD process) good lineages carry the history of their incipient species. An old lineage is indeed more likely to have a pool of incipient lineages (for instance, when *λ*_3_ is high, their number increases exponentially with time) and therefore less likely to go extinct. The equations we used for the probabilities of speciation 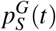 and extinction 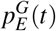 to the case of a good lineage with no incipient species, and are approximations otherwise. We expect that these approximations will affect mainly the extinction rate, as we ignore the buffering effect on extinction of incipient lineages that exist at the time when the rate is computed. The equivalent extinction rate could therefore overestimate extinction probabilities.

#### Speciation and extinction probabilities under the PBD

In order to compute 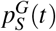 and 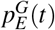 for any time *t* ∈ [0, *T*], we first need to compute probabilities associated with incipient lineages.

Starting from an incipient lineage, the two exclusive possible ways leading to extinction within a horizon *t* are (figure 2, upper left panel): (a) the first event to occur within the time horizon is an extinction or (b) the lineage forms two incipient lineages after a time *u*≤ *t* and both incipient lineages go extinct within the remaining time *t*− *u*. The probability for an incipient lineage and its potential descendants to go extinct within a time *t* thus satisfies the following equation:

**Figure 2.**
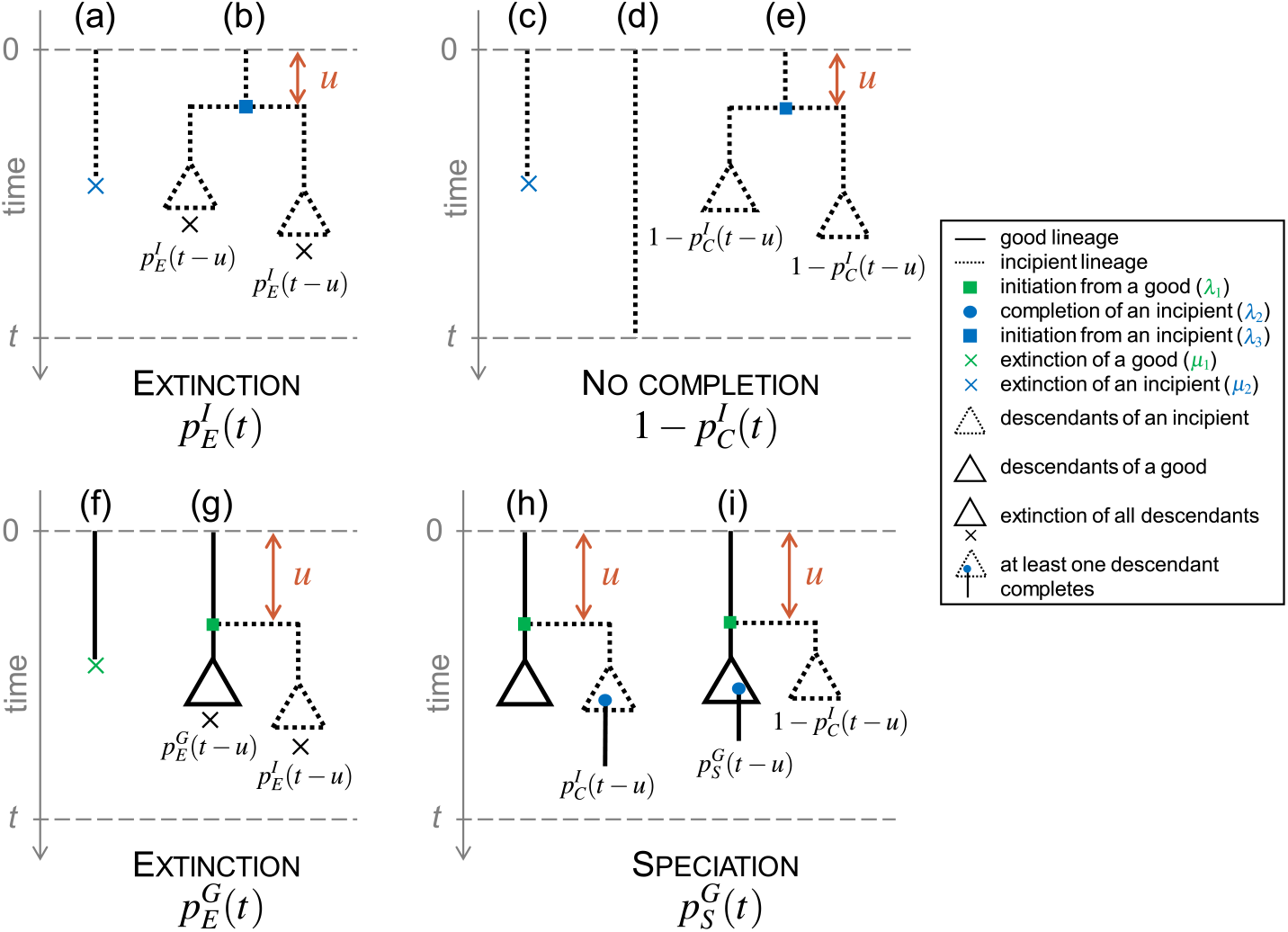
Alternative scenarios by which main outcomes occur in the protracted birthdeath model. Upper panels: starting from an incipient lineage at time 0, decomposition of the possible exclusive ways of extinction (left) and no completion (right) before a given time *t*. Bottom panels: starting from a good lineage at time 0, decomposition of the possible exclusive ways of extinction (left) and speciation (right) before a given time *t*. Dotted lines represent incipient lineages, solid lines represent good lineages. Triangles summarize the subtrees containing the potential descendants of an ancestor lineage, with the condition that all of them go extinct within the remaining time (indicated by a cross), or that one of them completes speciation (indicated by a blue dot), or that none of these events occur. The probabilities written under the triangles correspond to the probability of the described event.

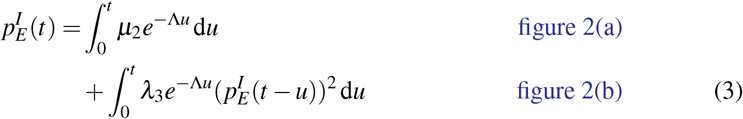

where Λ = *λ*_2_ + *λ*_3_ + *µ*_2_.

The solution to this integral equation is (appendix A1):

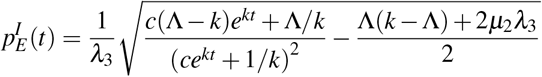

with 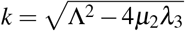 and 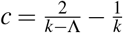.

Instead of 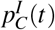, it is easier to calculate 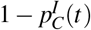, the probability of non-completion. Starting from an incipient lineage, the three exclusive ways not to complete speciation are (figure 2, upper right panel): (c) the first event to occur is the extinction of the lineage within a time *t*, (d) survival of the lineage during a time *t* without completion, extinction or initiation, or (e) initiation of speciation after a time *u*≤ *t* and non-completion of any of the daughter lineages within the remaining time *t* − *u*. The probability of speciation completion of an incipient lineage (or any of its potential descendant) within a time *t* thus satisfies the following equation:

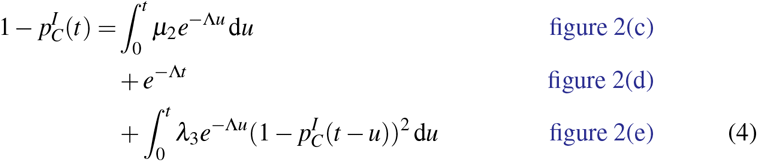

This equation can be solved numerically by solving an ordinary differential equation (appendix A2).

Starting now from a good lineage, the two exclusive possible ways leading to extinction as first event within a horizon *t* are (figure 2, bottom left panel): (f) the lineage directly goes extinct within a time *t* or (g) the lineage forms an incipient lineage and both daughter lineages (one good and one incipient) and their potential descendants die within a time *t* − *u* without speciation. The probability for a good lineage and its potential descendants to go extinct within a time *t* thus satisfies the following equation:

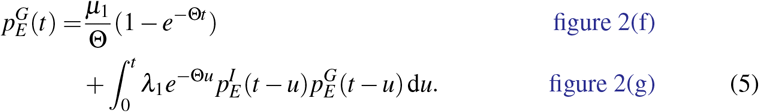

where Θ = *λ*_1_ + *µ*_1_.

This equation can be solved numerically, provided that we already have a solution for 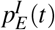 (appendix A3).

Starting from a good lineage, the two exclusive ways in which speciation occurs within a time *t* are (figure 2, bottom right panel): (h) the lineage initiates speciation after a time *u* ≤ *t*, and the incipient lineage completes speciation within the remaining time *t* − *u*, or (i) the lineage initiates speciation at a time *u*≤ *t*, the completion of this incipient lineage fails, and the good lineage speciates within the remaining time *t* − *u*. The probability that a good lineage fulfils speciation within a time *t* thus satisfies the following equation:

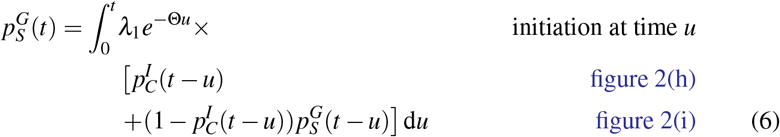

This equation can be solved numerically (appendix A4).

For all numerical integrations, we used the module `SciPy`in Python (Virtanen et al. 2020).

After solving these equations numerically and using equation 2, we obtain the timedependent equivalent birth rate 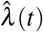 and death rate 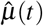.

### 2.3 Equivalent constant BD rates

Defined as above, the equivalent rates 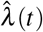 and 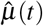 depend on time, even though the parameters of the PBD are constant. However, as we will show later and in agreement with previous results (Etienne and Rosindell 2012), the time-dependency is particularly manifest towards the present. Intuitively, given enough time, constant rates of initiation, completion and extinction result in constant equivalent speciation and extinction rates. However, towards the present (i.e., towards the end of the PDB process), incipient lineages did not have enough time to complete speciation, resulting in a decline in speciation rates. As one of our main goals here is to understand how speciation initiation, completion and extinction translate into macroevolutionary speciation and extinction rates, we now introduce equivalent constant BD rates, meant to represent the relationship between the parameters of the PBD and BD process far from the present.

We define equivalent constant BD rates, 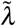 (birth rate) and 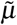 (death rate), as the parameters of the standard BD process such that the probability of speciation and the expected time to speciation match those under the PBD process with rates *λ*_1_, *λ*_2_, *λ*_3_, *µ*_1_, and *µ*_2_, starting from a good lineage. The probability of extinction does not bring any additional information since extinction and speciation events are complementary over infinite time. Intuitively, we expect that the equivalent time-dependent birth and death rates tend towards 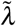 and 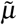 in the past; the equivalent time-dependent birth rate then declines toward the present.

We shown that equivalent constant-time rates are given by the following expressions (see details appendix A5):

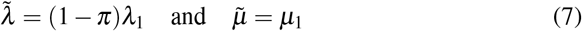

where 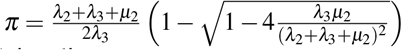 is the probability of non-completion of an incipient lineage.

Equation 7 shows that, far from the present, the equivalent death rate is exactly the rate of extinction of good species, and the equivalent birth rate is directly proportional to the rate of speciation initiation from a good species, with a coefficient of proportionality that represents the probability of completion without time horizon and depends on the rates specific to incipient lineages (initiation, completion and extinction). As in the case of the time-dependent equivalent rates, these rates correspond to the case of a process starting with a good lineage without incipient lineages, and the equivalent extinction rate may thus overestimate actual extinction probabilities under the PBD.

In order to better understand the influence of each parameter of the PBD process on the equivalent constant birth rate 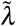, we calculate partial derivatives of this function with respect to the different parameters. A high partial derivative with respect to a given parameter reflects a strong influence of this parameter on the equivalent birth rate, and therefore that the corresponding step of the speciation process may be limiting. We compute the relative influence of a given parameter as the ratio between the absolute partial derivative with respect to this parameter and the sum of the absolute partial derivatives with respect to all other parameters. We perform these analyses both for the simplified PBD model where good and incipient lineages have the same rates of initiation (*λ*_1_ = *λ*_3_) and extinction (*µ*_1_ = *µ*_2_) and for the full PBD model. Detailed calculations are provided in appendix A6.

### 2.4 Simulations under the PBD process and equivalent BD processes

Although the equivalent rates were designed to construct BD processes that approach the PBD process, these processes are not identical. We used simulations to compare reconstructed trees (i.e., trees of extant species) generated by the PBD process and their equivalent BD processes. In each of these simulations, we consider the trees of extant good species, disregarding extinct and incipient lineages. We compared trees simulated under the constant-rate PBD model with trees simulated under the corresponding time constant (BD) and time-varying (varBD) equivalent BD models. We independently varied each of the 5 parameters of the PBD model, each taking 5 values (the default value and 2 above, 2 below, with amplitude chosen to guarantee computational tractability), resulting in 25 parameter combinations. For each combination, we computed the trajectories of equivalent time-dependent birth and death rates over 15 million years – approximated by piecewise-constant birth and death rates over 200 intervals – and simulated 500 tree replicates using the R library `TreeSim` (Stadler 2011). The values of these rates are given in supplementary figure S4. Each tree simulation was conditioned on the survival of two extant lineages (up to the failure of 100 simulation attempts for each replicate), starting from a single stem branch. We also generated the same number of trees under the PBD model for each combination of parameters using the `PBD` package (Etienne and Rosindell 2012). Finally, we simulated trees under the BD model with equivalent constant rates, using the package `TreeSim`. We expect these simulations to deviate the most from those obtained under the PBD process. We compared the outputs of the simulations under the three models in terms of species richness at present (SR), tree shape and tree topology.

To analyze tree shape, we used the *γ* statistic (Pybus and Harvey 2000), computed with the package `ape` (Paradis and Schliep 2019). The *γ* statistics quantifies the relative position of the internal nodes of a tree and compares it to the expectations under a pure-birth (Yule) model. *γ >* 0 corresponds to trees where internal nodes are closer to the tips than expected under Yule’s model, while *γ <* 0 corresponds to trees where internal nodes are closer to the root.

To analyze tree topology, we used the stairs2 balance index (Norström et al. 2012), computed with the package `treestats` (Janzen and Etienne 2024). This statistic measures the mean size ratio between the smaller and larger pending subtree for all vertices. Stairs2 is higher for trees with more balanced subtrees and lower for more imbalanced trees. The stairs2 statistic has been shown to perform well (Khurana et al. 2023; Kersting et al. 2024) and is less sensitive to tree size than other statistics such as Aldous’ *β* (Aldous 2001).

### 2.5 Tip speciation rate estimates

As we will show later and already mentioned, the equivalent birth rate declines close to the present, suggesting that speciation rates estimated at the tips of phylogenies may poorly reflect the underlying speciation process. To evaluate this effect, we computed the widely used DR (diversification rate) statistic (Jetz et al. 2012; Title and Rabosky 2019) at the tip of all the trees we simulated under the PBD process, using the package `epm` (Title et al. 2022). Next, for each tree, we compared the median DR over all extant leaves to the constant-rate equivalent birth rate. In order to evaluate deviations due solely to the use of DR (a biased estimator of speciation rate), we also computed DR at the tips of trees simulated under the constant-rate equivalent BD process.

### 2.6 Ability to recover equivalent constant BD rates by fitting the BD model to truncated trees

If we acknowledge that speciation in nature usually takes time, birth and death rates estimates obtained by fitting a constant-rate BD model to empirical reconstructed trees are hard to interpret. As noted above, under a constant rate PBD model, we expect equivalent birth rates to approach the equivalent constant birth rate 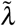 in the past, and to decline closer to the present. Hence, we can expect that speciation rate estimates obtained by fitting a BD model to the entire tree will have intermediate values below 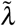. However, fitting a BD model to to older parts of the tree should provide good estimates of the equivalent constant BD rates.

To test this expectation, we truncated the phylogenies simulated under the PBD process in section 2.4 at different time points in the past, and fitted a constant BD model to these truncated phylogenies, using the dedicated function `fit bd in past` (Lewitus et al. 2018; Perez-Lamarque et al. 2022) from the R package `RPANDA` (Morlon et al. 2016). We fitted a constant-rate birth-death model to both entire trees and truncated trees “sliced” at 17 regularly spaced time points between the present and 4 million years before the present. Finally, we compared the speciation and extinction rate estimates obtained with various truncation times to the analytically-derived equivalent constant BD rates.

## 3. Results

### 3.1 Equivalent time-dependent BD rates

We used equations 1 and 2 and numerical solutions of the equations describing the probabilities of speciation, completion and extinction with time to derive equivalent time-dependent birth and death rates (thereafter simply referred to as birth and death rates) for a large range of parameter values.

We find that birth rates decrease close to the present, reaching 0 at present (time *t* →*T*, see figure 3). Death rates depend less on time and can be considered almost constant with time if we neglect a small decrease followed by a short increase when *t* →*T*. In the past (*t* →0), birth and death rates converge to the values predicted by our analytical expression of the equivalent constant BD rates (equation 7). This provides indirect (graphical) evidence that the constant equivalent birth-death rates can be considered as asymptotic rates of the time-dependent equivalent birth-death rates.

**Figure 3.**
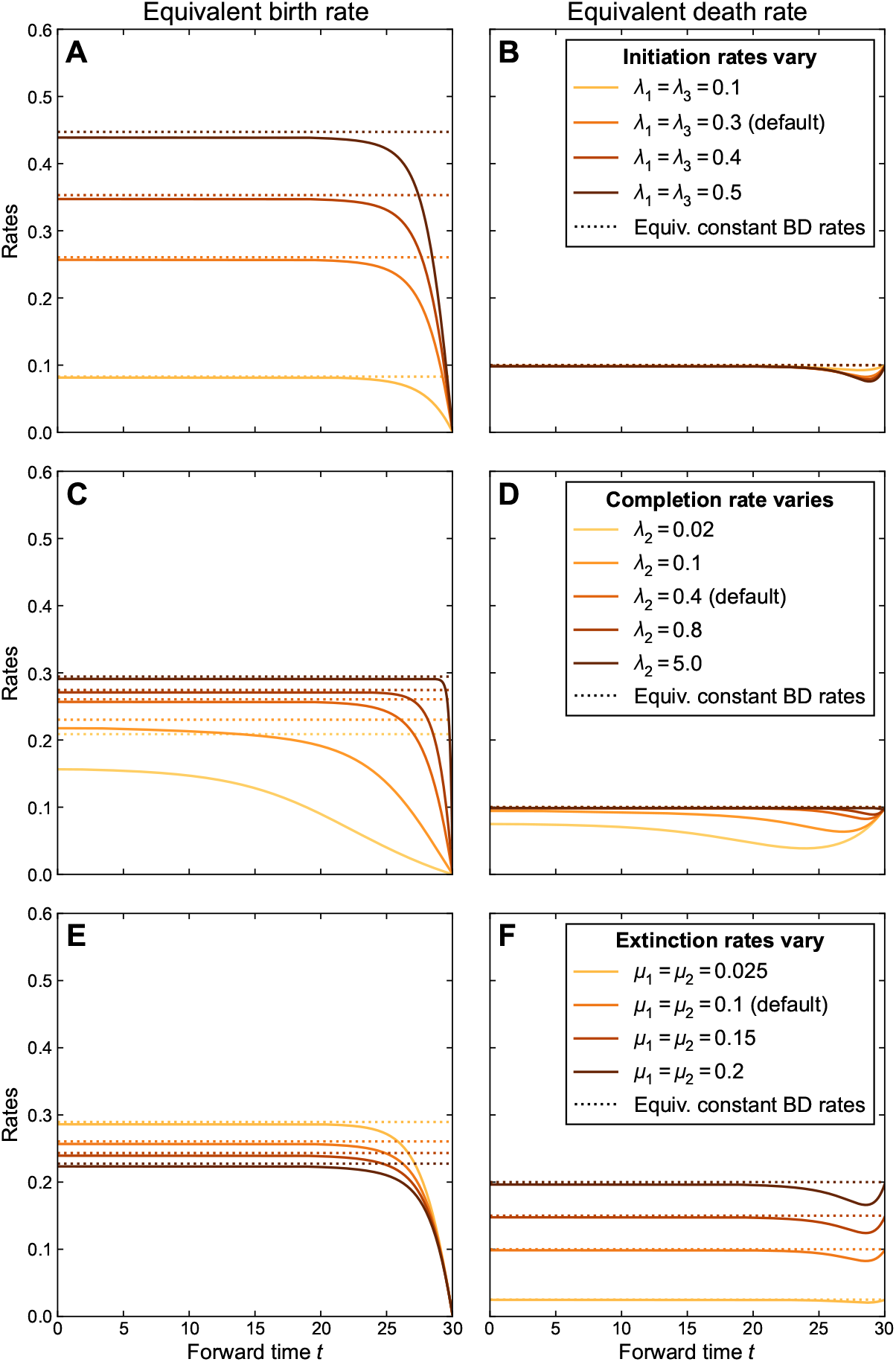
Influence of the parameters of the protracted birth-death (PBD) process on equivalent birth-death (BD) rates. Solid lines represent equivalent BD rates derived from equations 1 and 2 as a function of time for different values of PBD rates. Dotted lines represent constant equivalent BD rates derived analytically from equation 7 for the same PBD parameters. In the top/middle/bottom row the initiation/completion/extinction rates vary, with the other rates constant (default values are *λ*_1_ = *λ*_3_ = 0.3, *λ*_2_ = 0.4, *µ*_1_ = *µ*_2_ = 0.1). In figures S2 and S3, we calculated the same rates with the 5 parameters varying independently. *t* = 30 corresponds to the present, *t* = 0 to the past.

Initiation rates have a strong effect on the birth rate (figure 3, panel **A**). In the past, birth rates converge to values scaling with the initiation rate, all other rates being equal, as expected from equation 7. As expected, lower completion rates result in lower birth rates, and an effect of the protracted nature of speciation that extends further into the past (panel **C**). In the limit *λ*_2_ →∞, the model converges to a pure BD model with constant rates, except very close to the present. Indeed, with high completion rates, incipient lineages complete speciation very fast and with high probability, so speciation occurs as soon as an initiation event occurs. Finally, birth rates are lower when extinction rates are higher, except closer to the present where the effect of extinction diminishes (panel **E**). This effect is entirely due to the extinction of incipient lineages (*µ*_2_), as the extinction rate of good lineages (*µ*_1_) has virtually no effect on the birth rate (supplementary figure S2). The higher extinction rate of incipient lineages renders speciation less likely, as incipient species more often go extinct before completing speciation.

Death rates closely match extinction rates (figure 3, panel **F**), entirely due to the effect of the extinction rate of good lineages, as expected from equation 7; indeed, the extinction rate of incipient lineages has no effect on the death rate (supplementary figure S2).

### 3.2 Equivalent constant BD rates

As shown by equation 7 and already described above, the equivalent constant death rate equals the extinction rate of good species; the equivalent constant birth rate scales with speciation initiation, it increases with the completion rate and decreases with the extinction rate of incipient lineages (figure 3, dashed lines). We better characterized the influence of each step of the speciation process (i.e., initiation, survival of incipient species and completion) on 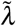, interpreted as the macroevolutionary speciation rate, by computing relative partial derivatives. This allows us to identify the steps of the PBD process that may limit macroevolutionary speciation rates figure 4. We identify that speciation initiation is limiting when its rate is low and the completion rate is high (regions 1 and 2). Speciation completion is limiting when its rate is low compared to the other parameters (regions 3 and 5). An increase in the population extinction rate has most effect when this rate is low (region 6) and when initiation rate is high compared to the completion rate (regions 4 and 5).

**Figure 4.**
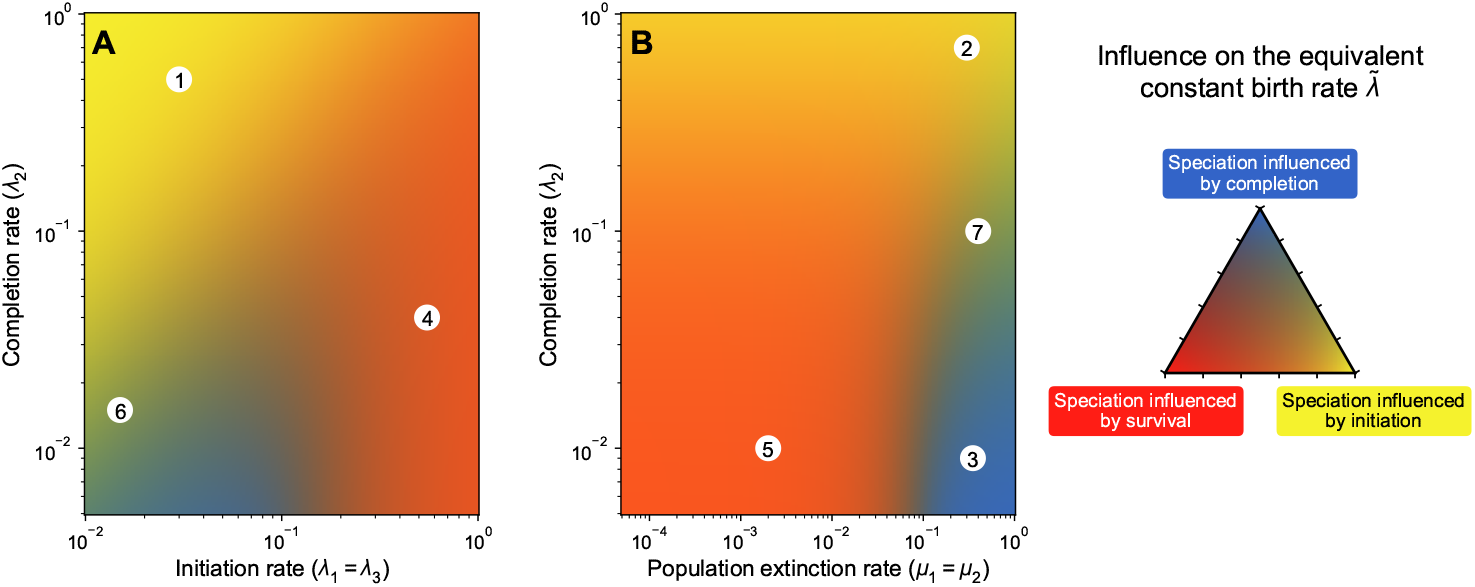
Relative influence of the parameters of the PBD model on the equivalent constant birth rate. Colors indicate which of the PBD process among initiation, completion and population extinction limits the equivalent constant birth rate 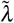 most, as a function of (**A**) the initiation and completion rates and (**B**) the population extinction and completion rates, with the color code explained on the triangle in the right. A yellow region (e.g. 1 or 2) indicates a combination of parameters where the most influential parameter on the birth rate is the rate of initiation. A blue region (e.g. 3) indicates a combination of parameters where the most influential parameter is the rate of completion *λ*_2_. An orange region (e.g. 4 or 5) indicates a combination of parameters where both population extinction and speciation initiation are influential. Green regions (e.g. 6 or 7) indicate situations where both speciation initiation and completion have a similar influence on the birth rate. In all cases, the rates of initiation and completion have a positive influence on the birth rate and the rate of population extinction has a negative influence. The detailed methods are explained in appendix A6 and the values of the relative influence are provided in supplementary figure S5. When they do not vary, default values of the parameters are 0.1.

### 3.3 Trees generated by the PBD process and equivalent BD processes

We compared the size, shape and topology of trees generated under the PBD model to those of trees generated under their equivalent time-constant and time-varying BD models (figure 5). As expected, tree size (SR) increases with higher rates of speciation initiation and completion, and decreases with higher rates of extinction of good and incipient lineages (figure 5, top row). These trends are well captured by both the equivalent time-varying and time-constant models, with the exception of the increase in species richness with the rate of initiation from incipient lineages *λ*_3_. Compared to the PBD model, the equivalent BD models produce larger trees when *λ*_3_ is small, and smaller trees when *λ*_3_ is large. *λ*_3_ has little influence on the size of trees generated under equivalent BD models, consistently with the weak influence of *λ*_3_ on the equivalent rates (see supplementary figure S3**C** and **D**).

**Figure 5.**
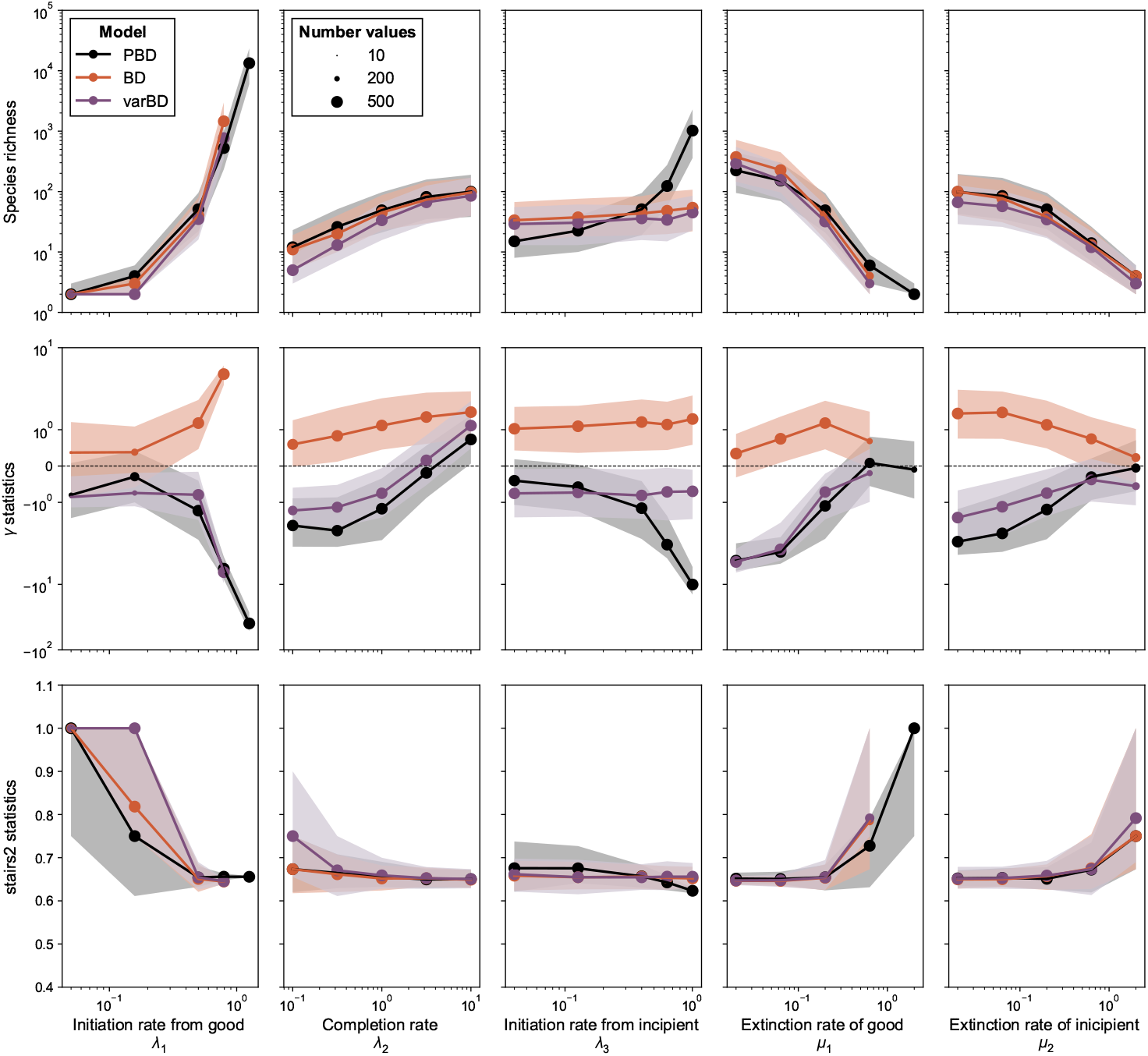
Statistics of trees generated under the protracted birth-death (PBD) process and its equivalent birth-death (BD) processes. By row: species richness SR, *γ* shape statistic and stairs2 balance index for trees generated under the three models (PBD, equivalent constant rate BD and equivalent time varying rate BD) for different values of the parameters of the PBD model. In each column, only one of the PBD parameters varies, with the others held constant (default *λ*_1_ = 0.5, *λ*_2_ = 1.0, *λ*_3_ = 0.4, *µ*_1_ = 0.2, *µ*_2_ = 0.2). For each set of parameters of the PBD model, BD and varBD trees were generated under equivalent birth and death rates computed using equations 1, 2 and 7. The line corresponds to the median of statistics across all 200 replicates, the shaded area indicates the first and last quartile of the statistics. The size of the dots indicates the number of valid data for which the statistics could be calculated.

As expected given unachieved speciation close to the present, the PBD process results in trees with negative *γ* values (reflecting long terminal branches), unless the completion or extinction rates are high (figure 5, middle row). *γ* decreases with increasing ratios of speciation initiation to speciation completion rates, i.e. when the protracted nature of the speciation process is more pronounced. The equivalent variable-rate BD model captures these trends relatively well, while the constant-rate BD model produces trees with positive *γ* values, indicating nodes closer to the tips. The drop towards 0 in the equivalent time-variable birth rates captures the shortage of speciation events close to the present induced by the protracted process, except when the rate of initiation from an incipient lineage (*λ*_3_) gets large. In this case the PBD process produces trees with increasingly long branch lengths, while the distribution of nodes generated under the equivalent time-dependent BD model remains stable.

Under most PBD parameters, stairs2 values of tree topology are stable, close to 0.65, and well reproduced by both the time-constant and the time-variable equivalent BD models. Higher values of stairs2 (reflecting more balanced trees) are observed in parameter ranges leading to small trees, and likely reflect tree size rather than a true difference in tree balance.

Deviations observed between trees simulated under the PBD process and those simulated under the equivalent time-varying BD model under some *λ*_3_ values likely come from the approximation we made when establishing the link between equivalent rates and the probabilities of speciation and extinction under the PBD (see section 2.2). If, as expected, the equivalent extinction rate overestimates extinction probabilities, this explains why we obtain smaller trees with less negative *γ* values, as extinction pushes nodes towards the present (the “pull of the present”; Nee et al. 1994). This is especially pronounced if *λ*_3_ is high; in this case, there are many incipient lineages that are not accounted for (the ones that originated before the time at which the rates are calculated).

### 3.4 Tip speciation rate estimates

DR tip speciation rates estimated on trees simulated under the PBD process are generally below equivalent constant birth rates (supplementary figure S8), as expected given the decrease in the equivalent birth rate near the present. This is not due to biases linked to the use of the DR statistic, as DR computed on trees generated under the equivalent BD process closely match equivalent birth rates.

### 3.5 Recovery of equivalent BD rates by fit to truncated PBD trees

Our expectation that fits of a constant rate BD model to the “old” part of trees generated under a PBD process would provide good estimates of the equivalent BD rates was verified (supplementary figures S7a and S7b). When fitting a constant rate BD model to the entire tree (i.e., when the truncation time is zero), the estimated speciation and extinction rates are well below the expected equivalent constant rates. However when truncation time increases, these estimates converge to the expected equivalent rates. In the case of the extinction rate, estimates remain slightly below the expected equivalent rates, supporting our intuition that equivalent extinction rates should overestimate actual extinction events. The convergence occurs with a truncation time relatively close to the present, consistent with the observed time of decline of the equivalent time-dependent BD rates. This recent truncation time appears optimal, as estimates are not as good when more of the tree is truncated, probably due to a loss of statistical power with decreasing data size.

## 4 Discussion

In this study we derived predictions, under the protracted birth-death (PBD) model of diversification, for “equivalent” birth and death rates, meant to represent macroevolutionary speciation and extinction rates. We showed that the equivalent rates – in particular the birth rate – vary when the process approaches the present, but can be considered constant when far from the present. Our analytical predictions of the rates in the past allowed us to explore the importance of each step of the speciation process (i.e. initiation, survival, completion) in modulating the macroevolutionary speciation rate. We found that the initiation and survival rates in general play a much larger role than the completion rate. In addition, we showed that the constant equivalent birth rate can be estimated by fitting a standard birth-death process on truncated reconstructed trees. This opens the possibility to relate estimates of macroevolutionary speciation rates, which have major consequences for the build up of diversity over geological time scales, to the microevolutionary processes that modulate speciation initiation, population survival and speciation completion.

Our equivalent rates are distinct from “congruent” rates, defined by Louca and Pennell 2020 as a combination of birth and death rates that result in the same likelihood on reconstructed trees. First, the equivalent BD process with variable rates is only an approximation of the PBD process (see the following paragraph), and therefore the equivalent BD model is not strictly speaking congruent to the PBD model. Second, we found a unique solution to our equations, demonstrating the unicity of equivalent rates: they are the only one that yield the same speciation and extinction probabilities at all time. Any other scenario in the same congruence class would have the same likelihood on reconstructed trees, but not the same speciation and extinction probabilities. An implication of this observation is that a time-varying birth-death scenario fitted on a reconstructed tree is not guaranteed to provide the equivalent rates. The criterion we chose to define equivalent rates is also distinct from the criterion used by Pannetier et al. 2021 to find a time-dependent BD model that approximates the diversity-dependent BD model. In the latter paper, the authors computed rates yielding the same expected diversitythrough-time (DTT). We expect our equivalent rates to produce the same DDT as that of the underlying PBD process, but we have not demonstrated that there are no other combinations of rates that also yield the same DDT.

As expected given previous results on the protracted birth-death process (Etienne and Rosindell 2012), we find that equivalent birth rates decline to zero as time approaches the present. Close to the present, speciation is less likely to occur because it requires a delay. The decay in equivalent birth rate starts earlier when the completion rate is lower. Trees simulated under the equivalent time-dependent BD model have similar characteristics to trees simulated under the PBD process, in terms of tree size and shape (figure 5), meaning that the equivalent BD process captures the dynamics of speciation and extinction induced by the protracted model relatively well. It is however important to remember that the PBD and equivalent BD processes are not entirely interchangeable. PBD is a process with memory, which induces an age dependency of speciation and extinction rates that is not captured by the equivalent BD process. For example “old” species are more likely to have accumulated incipient lineages, which buffer their extinction risk, but this cannot be captured by an age-independent BD process. More generally, the approximations made in our equivalent BD process lead to an overestimation of the frequency of extinction events occurring under the PBD process, explaining some of the deviations observed when comparing simulated trees, in particular for high values of the rate of initiation from incipient lineages (*λ*_3_).

Before equivalent speciation rates start to drop, they can be considered virtually constant. In this regime, we derived analytical relationships between the equivalent birth and death rates and the parameters of the PBD process. These relationships show that we can expect macroevolutionary extinction rates to provide a relatively good approximation of the rate at which species go extinct, except if the rate of initiation from incipient lineages is large. They also show that macroevolutionary speciation rates are directly proportional to speciation initiation rates, with a coefficient of proportionality that depends on the rates of completion, initiation and extinction of incipient lineages. Hence, all aspects of the speciation process play a role in modulating macroevolutionary speciation rates, but some play a larger role than others, and which ones will likely depend on the ecology, genetics and biogeography of the species group considered. Indeed, by identifying regimes within which each aspect of the speciation process is expected to be the most influential (figure 4), we found that the rates of speciation initiation and population extinction often are the limiting factors. Completion rates are only limiting when they are very small and at intermediate values of speciation initiation rates, or high population extinction rates. Hence, a species that accumulates fast reproductive isolation will be characterized by a higher completion rate but this will not necessarily have a strong effect on the speciation rate. In their analysis of Australian rainbow skinks, Hua et al. 2022 estimated rates of speciation initiation and completion of 0.27 and 0.31, respectively, with low extinction rates. In this area of parameter space, speciation rates are clearly not limited by the speed at which reproductive isolation is acquired (relative influence ∼ 10^−7^), but more by the initiation of speciation (68%) and the persistence of lineages (32%) (supplementary figure S9).

Our results have implications for the phylogenetic analysis of diversification. Indeed, ignoring the fact that speciation takes time by fitting standard BD models to empirical phylogenies necessarily leads to model misspecification (e.g. when fitting a constant rate BD model) or misinterpretation (e.g. when interpreting the effect of protraction as a speciation rate decline). In the recent years, there has been a surge in the use of tip-rate speciation estimates, i.e. estimates of speciation rates at present across the tips of a phylogeny, obtained with statistics such as the diversification rate (DR; Jetz et al. 2012) statistic, or models such as the Bayesian analysis of macroevolutionary mixtures (BAMM; Rabosky 2014), the cladogenetic diversification shift (ClaDS; Maliet et al. 2019; Maliet and Morlon 2021), or the birth-death diffusion (BDD; Quintero et al. 2024). Future work investigating how speciation rates estimated with these methods on trees generated under the PBD process compare with equivalent rates will be useful. We conducted this exercise here with the DR statistic, and found that the obtained rates generally underestimate the equivalent constant birth rate, reflecting the convergence to zero of the equivalent birth rate close to the present. Hence, tip rate estimates are particularly challenging to interpret in terms of the underlying speciation process. Speciation rates estimated on deeper parts of the phylogeny approximate equivalent rates quite well, suggesting that these estimates indeed reflect the macroevolutionary outcome of the combined processes of initiation, survival and completion. Ideally, we would need to account for the protracted nature of speciation in every model used to infer diversification rates from phylogenies. By incorporating a protracted process in the State-dependent Speciation-Extinction (SEE) framework, Hua et al. 2022 made progress in this direction. A workaround is to truncate phylogenies, a method that has previously been used in another context (Phillimore and Price 2008). However, when truncating actual species splits to avoid the effect of protraction close to the present, this requires employing inference methods that account for this truncation. These have been implemented for only a limited set of models, such as the constant-rate, time-dependent and environment-dependent models developed in `RPANDA` (Morlon et al. 2016). More systematically implementing this truncation option in diversification models would be useful. Our tests indicate that truncating phylogenetic trees at intermediate time points yields the most accurate estimates of equivalent rates. As the optimal truncation time will depend on the extent to which speciation is protracted (i.e., the completion rate), we recommend to apply truncation at various time points and choose the time at which the estimated rates reach a plateau.

Our results also have implications for the ongoing effort to relate macroevolutionary speciation rates to microevolutionary processes (Rolland et al. 2023; Harvey et al. 2019; Rabosky 2016; Morlon et al. 2024). Indeed, microevolutionary processes act individually on each step of the speciation process (initiation, survival, completion), and our expressions of equivalent rates quantify how these combine to modulate macroevolutionary speciation rates. For example, the apparent decoupling between tip rate estimates of speciation and the speed at which reproductive isolation is acquired (Rabosky and Matute 2013; Freeman et al. 2022) is not surprising given our expectation that completion rates have a limited effect on the macroevolutionary speciation rate (and the limitations of tip rate estimates mentioned before). Etienne et al. 2014 observed a weak correlation between speciation completion rates estimated with the PDB model and speciation rates estimated with a BD model, on both simulated and bird phylogenies, especially when the completion rate was low. The authors interpreted this result as the limits of estimating speciation rates with a BD process when speciation is protracted. Our results suggest that the decoupling is in fact real and expected. We would expect the correlation to be especially weak when the completion rate is high rather than low, and the discrepency between these expectations and Etienne et al.’s observations could be linked to the limits of estimating speciation rates with a BD process. We, however, expect to find a correlation between macroevolutionary speciation rates and the rate of population formation, which reflects speciation initiation. This is indeed what was found by Harvey et al. 2017; the lack of correlation in other studies (Singhal et al. 2022; Singhal et al. 2018; Burbrink et al. 2023) may reflect cases when speciation is modulated by the rate of population extinction rather than by the rate of speciation initiation, as expected if the initiation rate is high and/or the population extinction rate is low.

By taking into account the time it takes to complete speciation, protracted birth-death models provide more biologically realistic models than standard birth-death models, and a theoretical framework for understanding how each step of the speciation process influences macroevolutionary speciation rates. A more systematic account of the protracted nature of speciation in phylogenetic analyses of diversification would both improve our estimates of diversification rates and our understanding of how microevolutionary processes combine to modulate macroevolutionary speciation rates.

## Code availability

The scripts used to make these analyses are available on https://github.com/pierre-veron/PBD_analog.

## Aknowledgement

We thank Thibault Juillard, Amandine Véber and Amaury Lambert for mathematical advice, and Beno ît Perez-Lamarque who kindly shared his code to estimate the BD rates on truncated phylogenies.

We also thank Tanja Stadler, Matt Pennell and Rampal Etienne for their comments of a previous version of the manuscript.

## Appendices

### A1 Resolution of the probability of extinction of an incipient lineage

In equation 3, let us do the variable change *v* = *t* − *u* to get rid of the variable *t* in the integral:

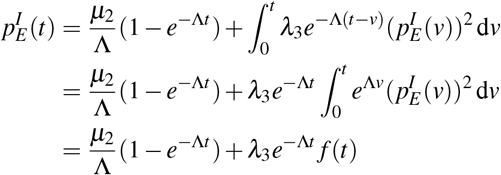

with *f* defined with, for all *t* ≥ 0:

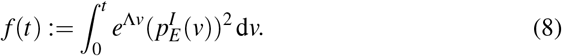

We note that

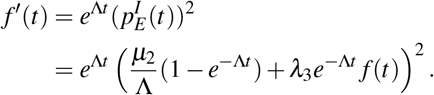

This is a non-linear ordinary differential equation (ODE) of first order, with the initial condition *f* (0) = 0.

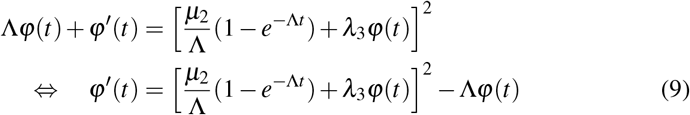

Let us check that the function

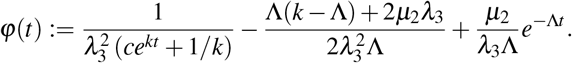

with 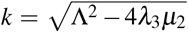 and *c >* 0 satisfies this last ODE. Inside the bracket of the right-hand side (RHS) of equation 9 we notice the simplification:

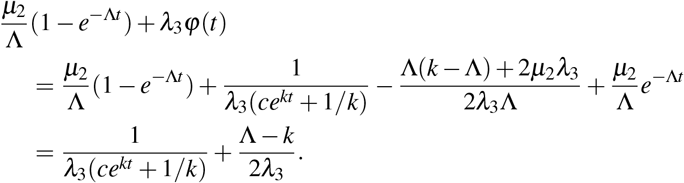

So the RHS of equation 9 is:

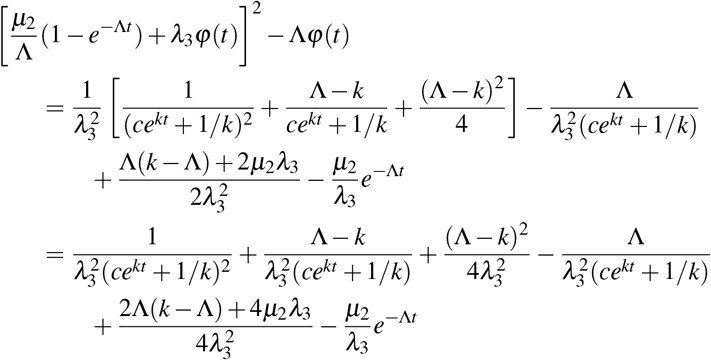

and since (Λ− *k*)^2^ + 2Λ(*k*−Λ) + 4*µ*_2_*λ*_3_ = *k*^2^ −Λ^2^ + 4*µ*_2_*λ*_3_ = 0 given the definition of *k*, this simplifies into:

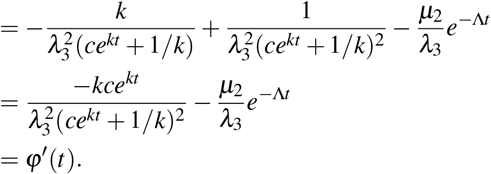

so the suggested φ is a solution of the ODE equation 9. The condition φ(0) = 0 imposes 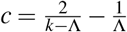. Finally, we have the solution of the original ODE:

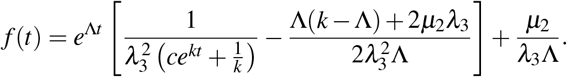

From equation 8 we have

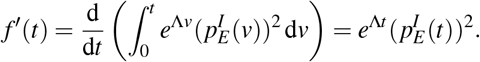

So the probability of extinction 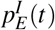 can be retrieved

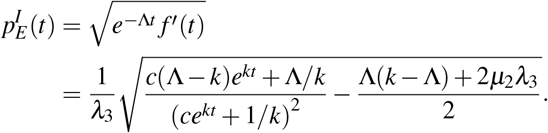

#### A2 Resolution of the probability of completion of an incipient lineage

To get rid of the variable *t* in the integral of equation 4, we do the change of variable *v* = *t* −*u*. This gives us:

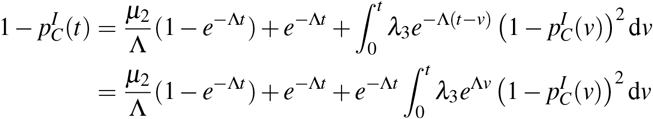

we now do a logarithmic change of variable in the integral, *z* = *e*^Λ*v*^:

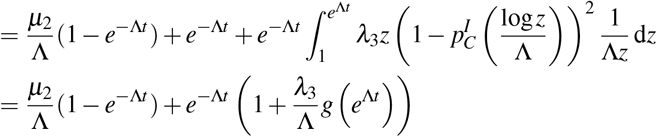

with, for *x* ≥ 1:

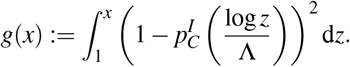

Noting that 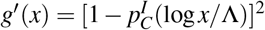, we have:

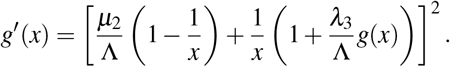

This is a non-linear ODE. With the initial condition *g*(1) = 0, it can be solved numerically and we retrieve the probability of completion of an incipient lineage with

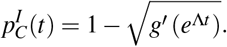

#### A3 Resolution of the probability of extinction of a good lineage

From equation 5 we do the same change of variable as in the previous part *v* = *t* −*u*:

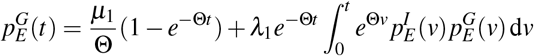

then with *z* = *e*^Θ*v*^ we have:

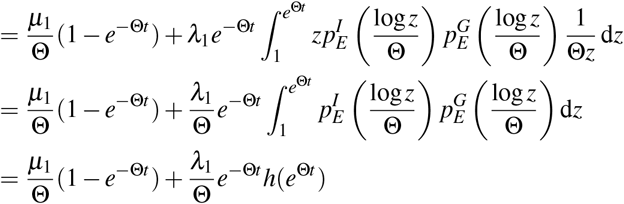

with, for *x* ≥ 1

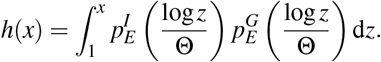

We note that

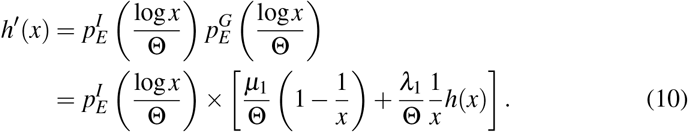

With the condition *h*(1) = 0, equation 10 gives the ODE satisfied by *h*. If we solve this equation numerically we can retrieve 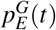:

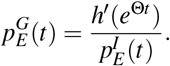

#### A4 Resolution of the probability of speciation of a good lineage

From equation 6, we do the same change of variable *v* = *t* −*u*, then *z* = *e*^Θ*v*^:

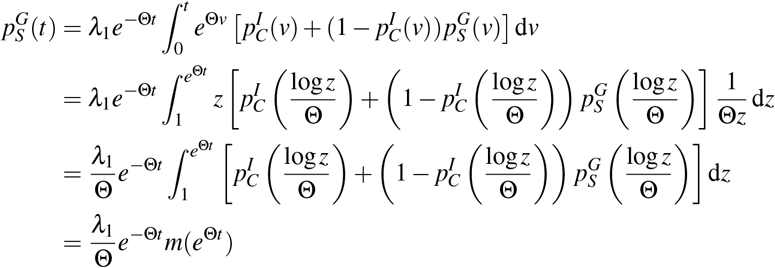

with, for *x* ≥ 0

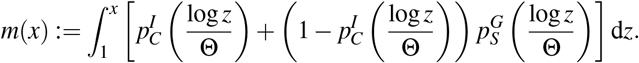

We note that

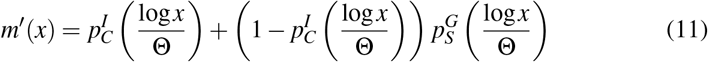

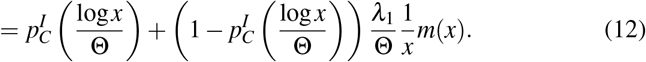

With the initial condition *m*(1) = 0, the equation 12 gives the ODE satisfied by *m*. If we solve this equation numerically we can retrieve 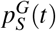 with the derivative of the solution *m*, from equation 11:

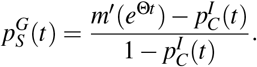

We show an example of the obtained numerical solutions of 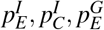 and 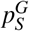 and empirical probabilities from simulations in supplementary figure S1.

#### A5 Calculation of the time-constant BD rates under the PBD model

##### Speciation probability and expected time for speciation under the BD model

Under the BD model with parameters *λ* and *µ*, the probability of speciation of a given lineage is the probability that a birth event occurs before a death event, i.e.

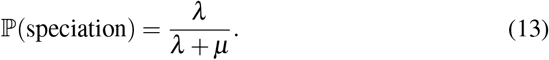

Conditionally on speciation, the expected time *T* it takes for a given lineage to speciate is

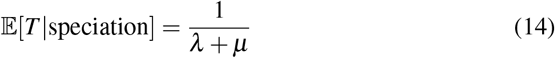

since *T* has the same distribution as min(*X,Y*) with 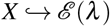 and 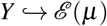 two independent clock variables representing speciation and extinction events. The minimum of two exponential independent processes is distributed exponentially with a parameter equal to the sum of the rates.

##### Speciation probability and expected time for speciation under the PBD model

Under the PBD model, a good lineage speciates if it generates at least one incipient lineage and one of these incipient lineages or one of the descendants of these lineages completes speciation.

Considering an incipient lineage *L* that has still no descendant, there are two outcomes: (*i*) the lineage or one of its descendants at least complete speciation (we will denote this event as “complete”), or (*ii*) neither the lineage nor any of its descendants complete speciation before dying (we will denote this event as “does not complete”).

Let us denote *π* := ℙ (*L* does not not complete) and *N* the number of direct descendants of the lineage before it dies or completes speciation, and *L* and *L*_1_, …, *L*_*N*_ those lineages. All the possible events that can happen to *L* (completion at rate *λ*_2_, new incipient lineage at rate *λ*_3_ or extinction at rate *µ*_2_) are independent point process. The first event to happen is therefore a point process with rate *λ*_2_ + *λ*_3_ + *µ*_2_ and the probability that it is the formation of an incipient lineage is *λ*_3_*/*(*λ*_2_ + *λ*_3_ + *µ*_2_). We can thus decompose:

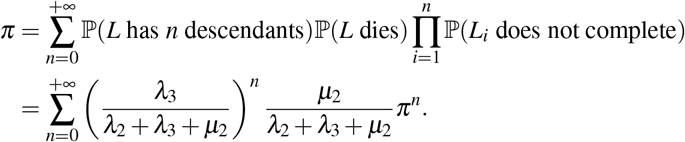

Therefore

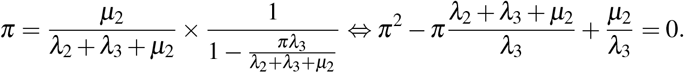

The solution of this equation in [0, 1] is

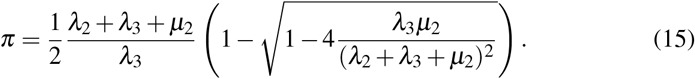

Let us now consider a good lineage. Ignoring the proportion *π* of incipient daughter lineages that will not complete speciation, we can consider the filtered point process with rates of successful initiation (1− *π*)*λ*_1_ and extinction *µ*_1_. Therefore the probability of speciation is the probability that a successful initiation occurs before the extinction, given by:

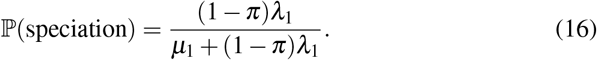

The speciation time is the branching time of the first successful incipient lineage to speciate. Therefore the speciation time is distributed as the result of the filtered point process and has the expected value:

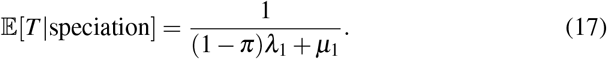

##### Expression of the equivalent rates of speciation and extinction

The equivalent rates of BD are the rates such that ℙ (speciation) and 𝔼 [*T*|speciation] are equal in both models. To do so, we set 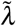 and 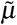 such that the pairs of expressions in equation 13/16 and 14/17 are equal. This gives us:

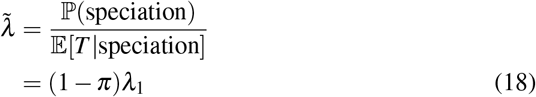

and

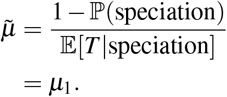

#### A6 Analysis of parameters limiting the equivalent birth rate

We measure the influence of the variation in each of the parameters of the PBD model on the equivalent birth rate 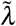. Parameters that influence 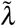 the most can be considered potentially limiting, as they can drive 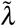 down depending on their values. We calculated the partial derivatives of 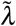, given by the expression equation 18, with respect to the PBD parameters *λ*_1_, *λ*_2_, *λ*_3_ and *µ*_2_ (not with respect to *µ*_1_, denoting the rate of extinction of a good lineage, because it has no influence on the equivalent birth rate).

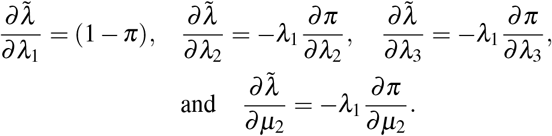

In what follows, we explicit the expressions (*i, ii* and *iii*) of the three partial derivatives of *π* with respect to the three parameters of the incipient lineages. From equation 15 we 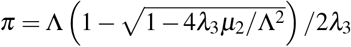 with Λ = *λ*_2_ + *λ*_3_ + *µ*_2_.

(i) 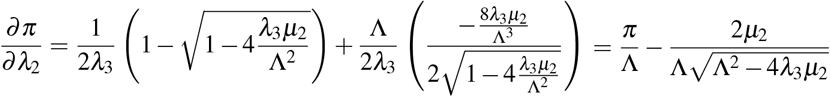

(ii) 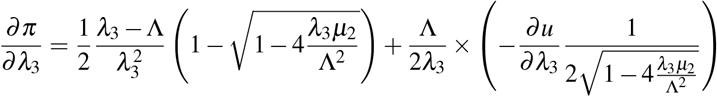

with *u* defined as 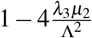, so 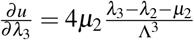, hence

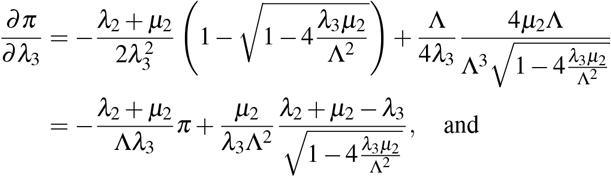

and

(iii) 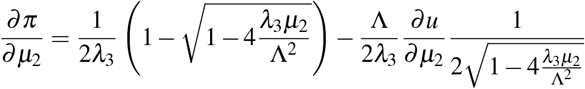

with 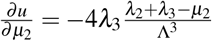

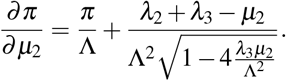

We then use the simplified framework of the PBD model where all rates of initiation are equal (*b* := *λ*_1_ = *λ*_3_) and all rates of population extinction are equal (*e* := *µ*_1_ = *µ*_2_). By the chain rule:

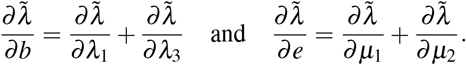

We then evaluate the relative influence of a parameter as the ratio between the absolute partial derivatives. For instance the relative influence of the rate of initiation *b* is:

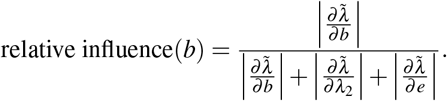

We provided a detailed plot of the values of this relative influence of the three simplified parameters in supplementary figure S5. The summary of these relative influences is provided on figure 4. A similar analysis was also done with the detailed model (with *λ*_1_≠*λ*_3_ and *µ*_1_≠*µ*2 a priori, supplementary figure S6).

