## Supplementary figures for "Speciation completion rates have limited impact on macroevolutionary diversification"

September 9, 2024

### Supplementary material

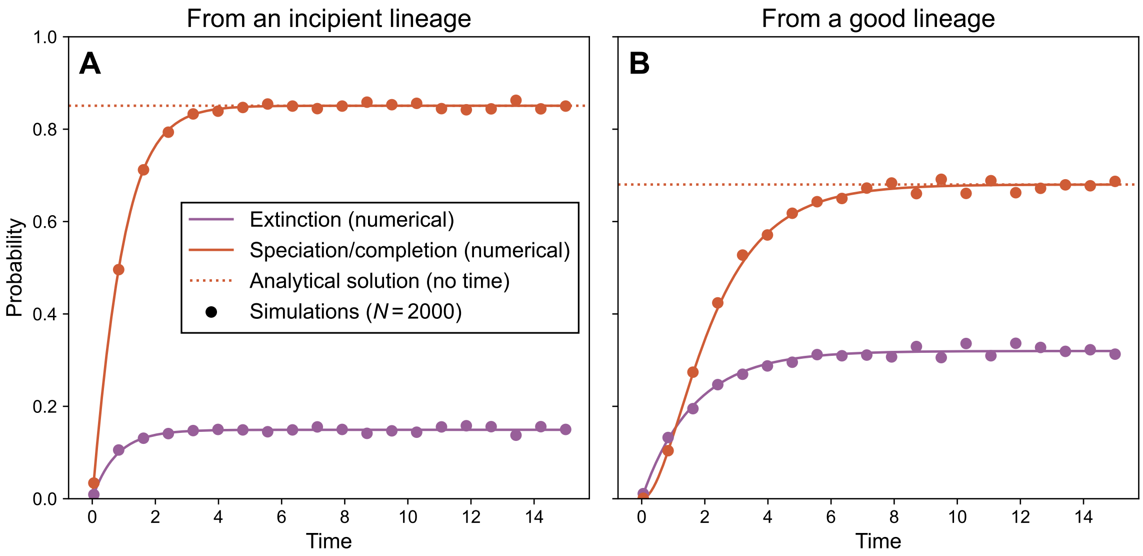


**Figure S1. Completion or speciation and extinction probabilities within a time horizon $t$ calculated numerically** (solid lines, see main text and appendix) **or empirically estimated on simulations** (dots). Panel **A** shows the probabilities starting with an incipient lineage ($p_E^I$, $p_C^I$) and panel **B** shows the probabilities starting with a good lineage ($p_E^G$, $p_S^G$). Each dot corresponds to $N = 2000$ replicates (each one starts with one lineage and runs for a finite time $t$) and shows the frequency of replicates ending by a speciation or extinction event within a time horizon $t$. The dotted line corresponds to the analytical expression of the speciation probability (equation 15 for an incipient lineage and equation 16 for a good lineage, in the main document). Parameters used: $\lambda_1 = 0.5, \lambda_2 = 0.8, \lambda_3 = 0.4, \mu_1 = \mu_2 = 0.2$, same notations as in Etienne and Rosindell 2012 and in the main document.

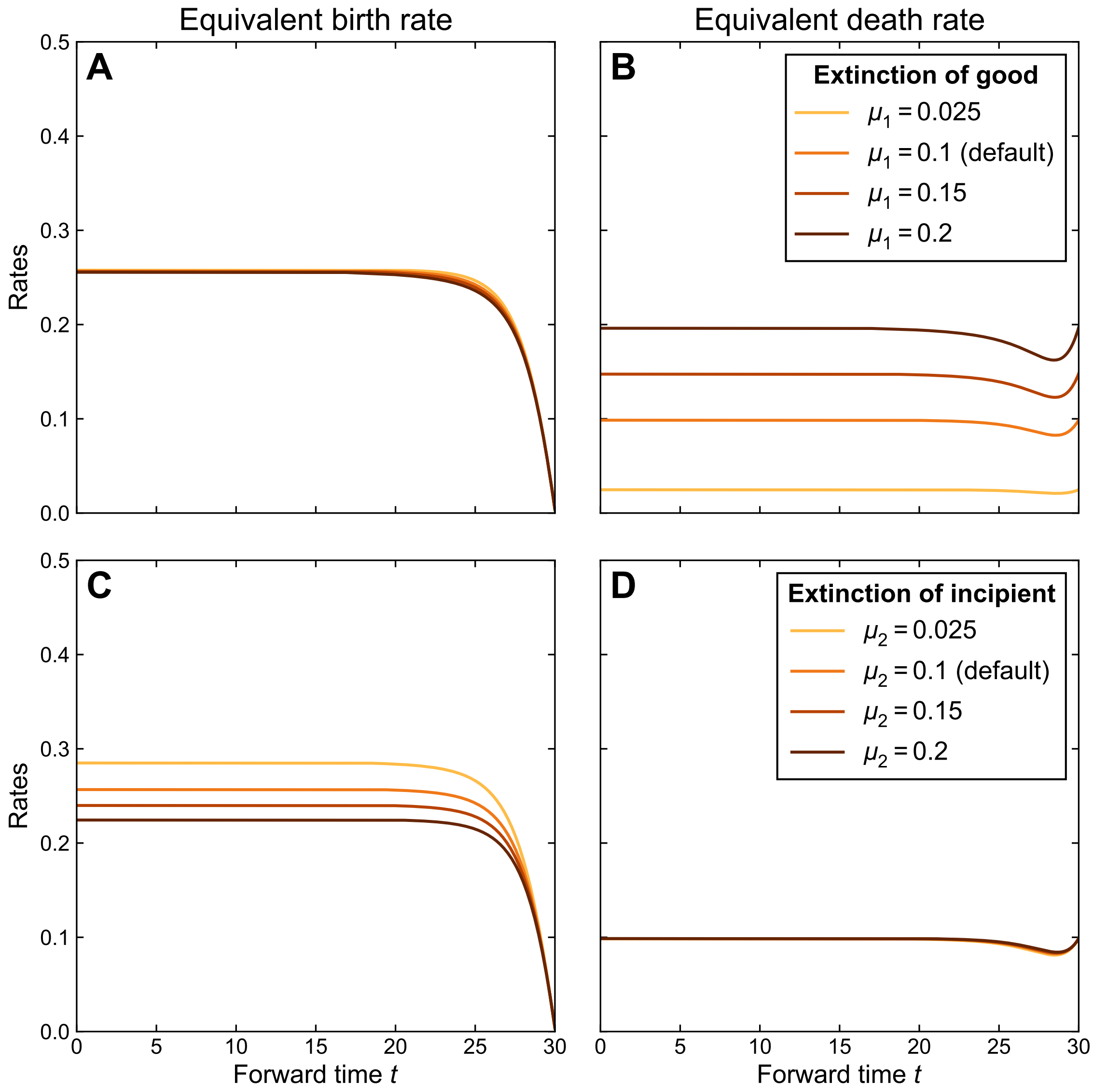


**Figure S2. Values of the time-dependent birth-death rates** derived from equations 1 and 2 (main document) as a function of time **as both extinction rates of the PBD model $\mu_1$ and $\mu_2$ vary independently**. In the top row (**A** and **B**) the extinction rate of good lineages varies while in the bottom row (**C** and **D**) the extinction rate of incipient lineages varies. When one rate varies, the other four rates are kept constant (default values are $\lambda_1 = \lambda_3 = 0.3, \lambda_2 = 0.4, \mu_1 = 0.1, \mu_2 = 0.1$).

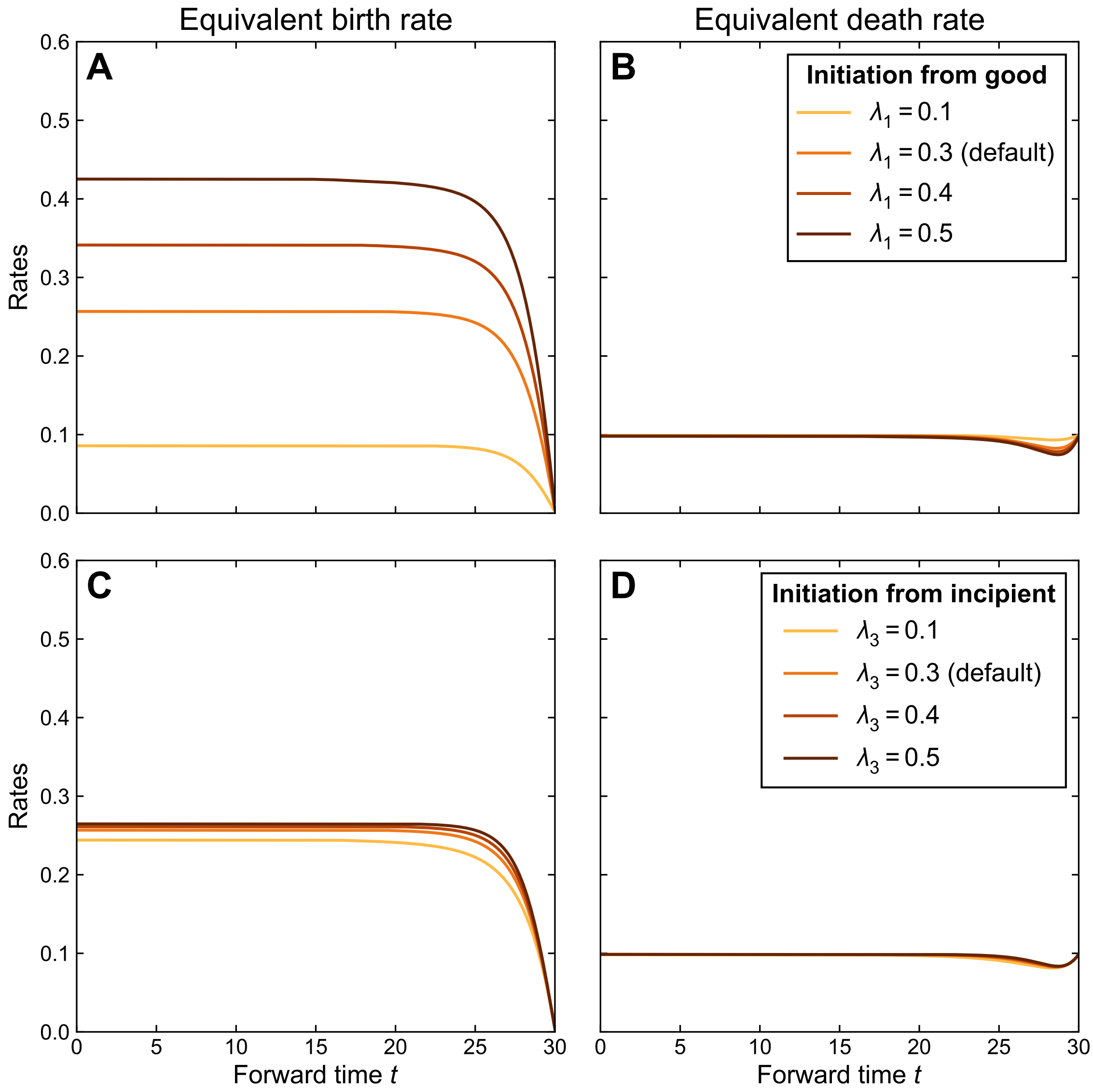


**Figure S3. Values of the time-dependent birth-death rates** derived from equations 1 and 2 (main document) as a function of time **as both initiation rates of the protracted birth-death model ($\lambda_1$ and $\lambda_3$) vary independently**. In the top row (**A** and **B**) the initiation rate from good lineages varies while in the bottom row (**C** and **D**) the initiation rate from incipient lineages varies. When one rate varies, the other four rates are kept constant (default values are $\lambda_1 = 0.3, \lambda_3 = 0.3, \lambda_2 = 0.4, \mu_1 = \mu_2 = 0.1$).

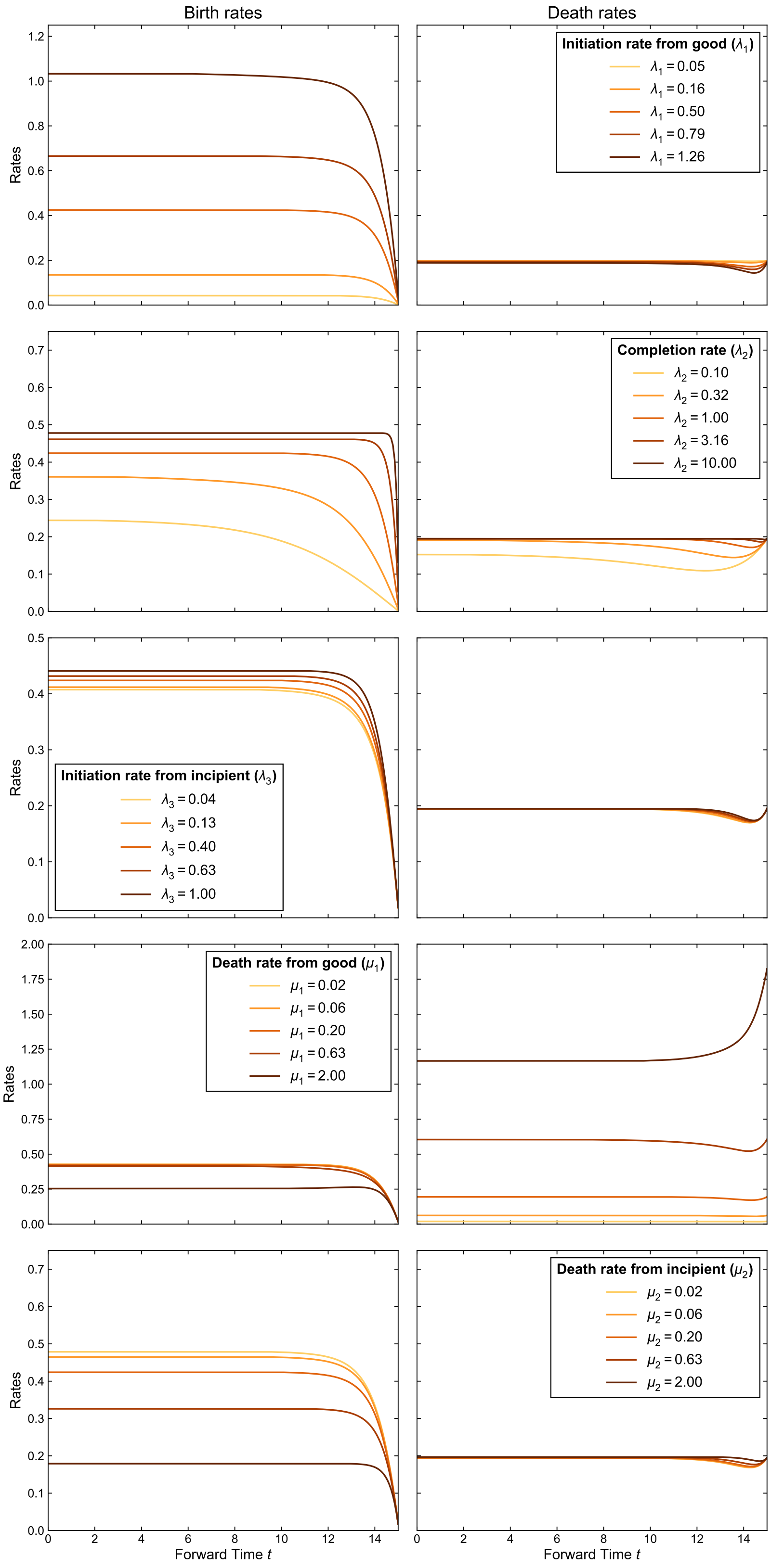


**Figure S4. Values of the time-dependent equivalent birth-death (BD) rates used in the simulations**. On each row, only one parameter of the protracted birth-death (PBD) model varies. The left column shows the birth rate and the right column shows the death rate. Each curve corresponds to a set of parameters of the PBD model. The resulting equivalent BD rates are used to generate trees under a time-dependant BD model that are summarized in figure 5 (main document).

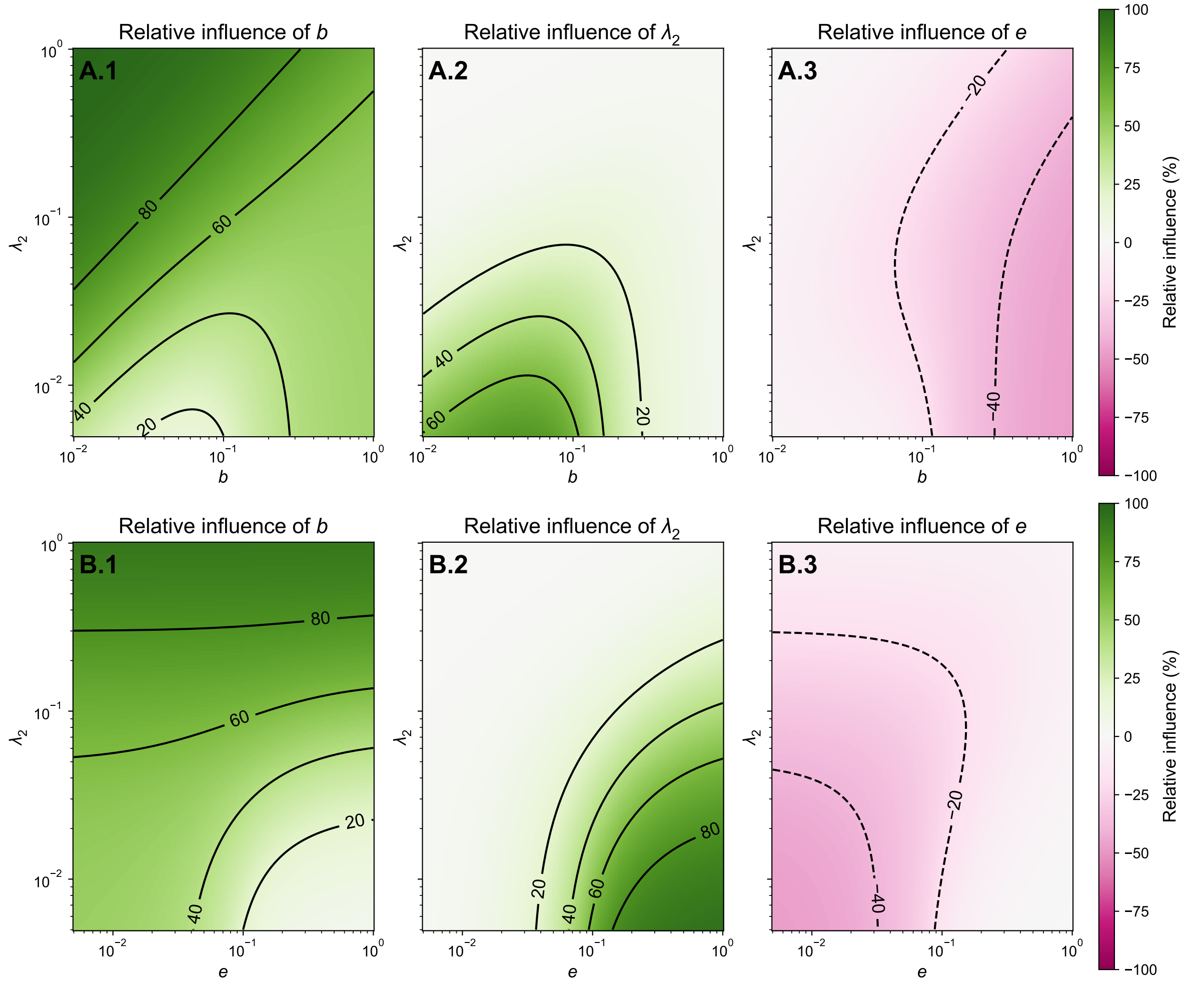


**Figure S5. Relative influence of the parameters of the *simplified* protracted birth-death (PBD) model** ($b := \lambda_1 = \lambda_3$ and $e := \mu_1 = \mu_2$) **on the constant equivalent birth rate** $\tilde{\lambda}$, as defined in appendix A6, as a function of $(b, \lambda_2)$ (panels **A**) and $(e, \lambda_2)$ (panels **B**). When unspecified, the values of the rates of the PBD model are taken at 0.1. The values shown here are summarized in figure 4 (main document).

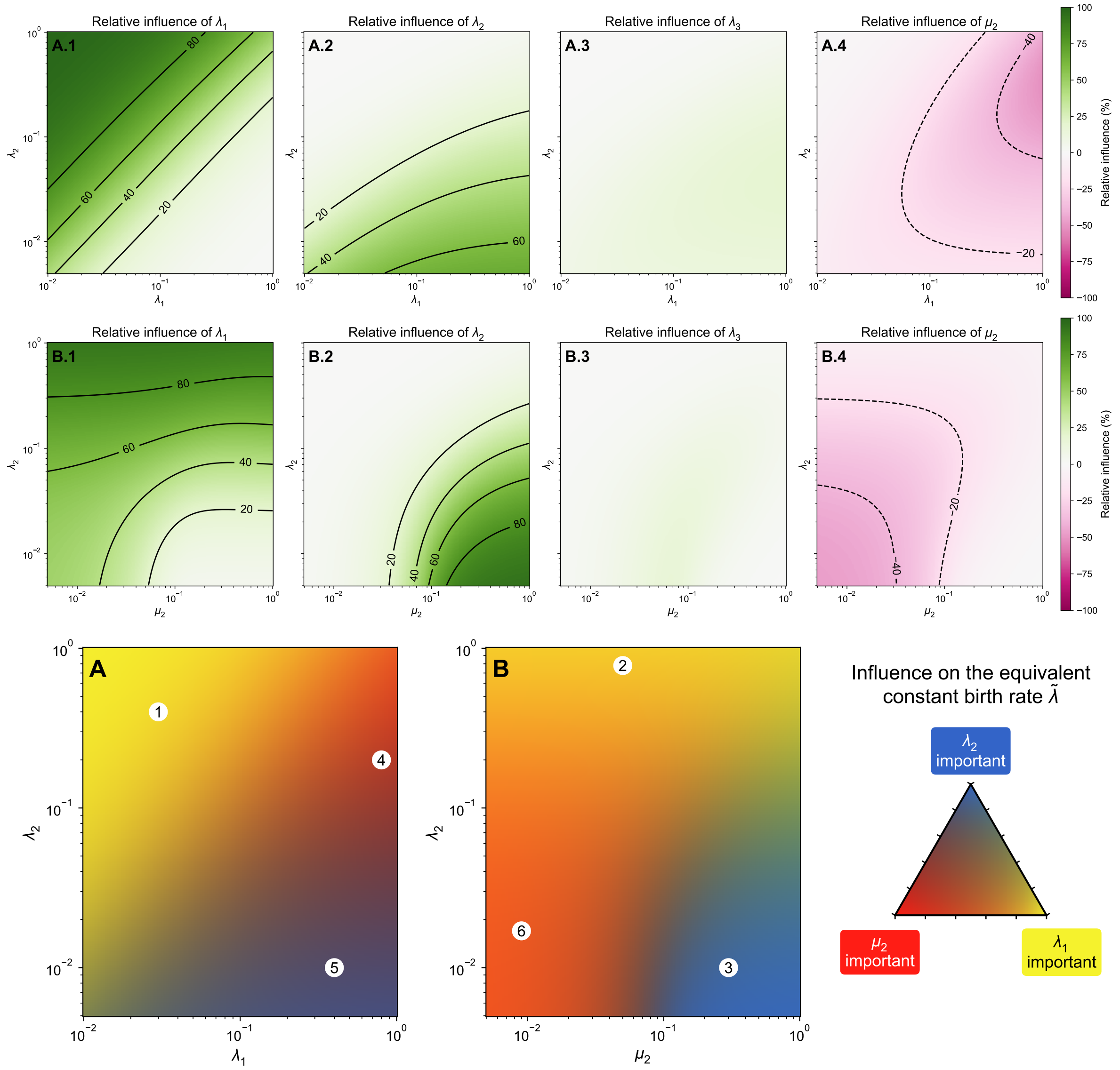


**Figure S6. Relative influence of the parameters of the PBD model on the constant equivalent birth rate** $\tilde{\lambda}$, as defined in appendix A6, as a function of $(\lambda_1, \lambda_2)$ (panels **A**) and $(\mu_2, \lambda_2)$ (panels **B**). When unspecified, the values of the rates of the protracted birth-death model are set at 0.1.

In the lower plot, colors indicate which of the parameters among $\lambda_1, \lambda_2$ and $\mu_2$ are most limiting in the variation of the equivalent constant birth rate $\tilde{\lambda}$, with the color code explained in the triangle on the right. A yellow region (e.g. 1 or 2) indicates a combination of parameters in which the most influential parameter on the birth rate is the rate of initiation $\lambda_1$. A blue region (e.g. 3) indicates a combination of parameters for which the most influential parameter is the rate of completion $\lambda_2$. A red region (e.g. 4) indicates a combination of parameters where the most influential parameter is the rate of extinction of incipient lineages $\mu_2$. Purple regions (e.g. 5), orange regions (e.g. 6) or green regions indicate situations in which several parameters have a comparable influences on the birth rate. In all cases, the influences of $\lambda_1$ and $\lambda_2$ are positive and the influence of $\mu_2$ is negative.

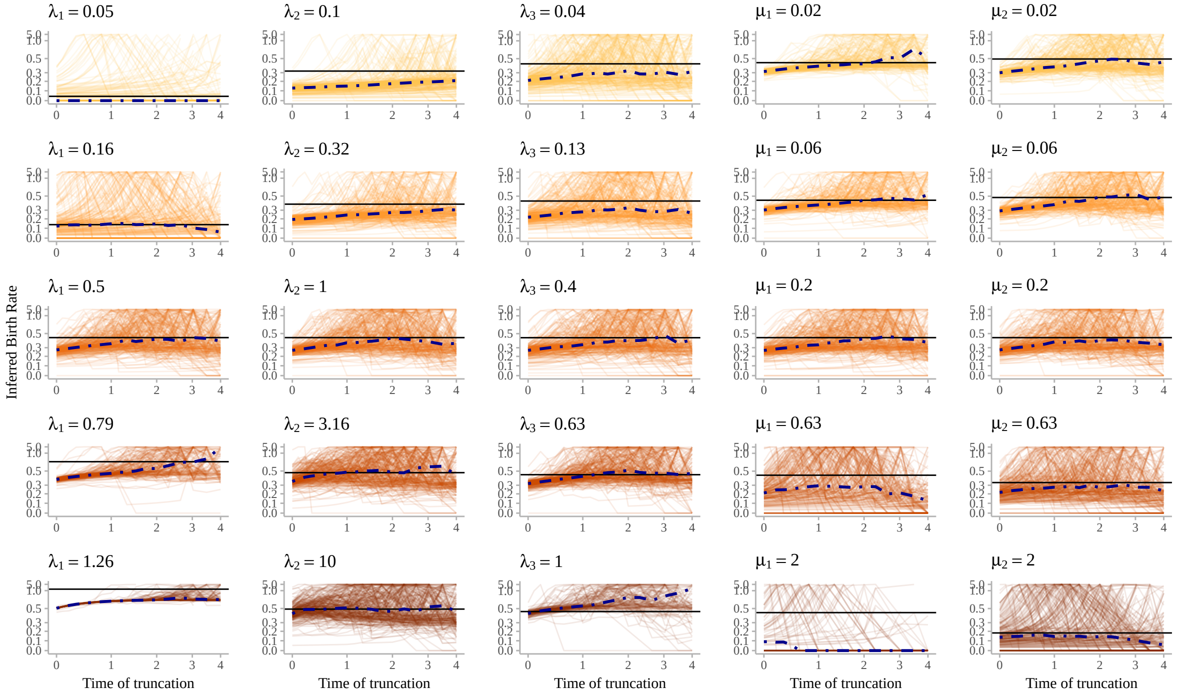


**(a)** Inferred birth rates ($\lambda$) across truncation lengths and replicates.

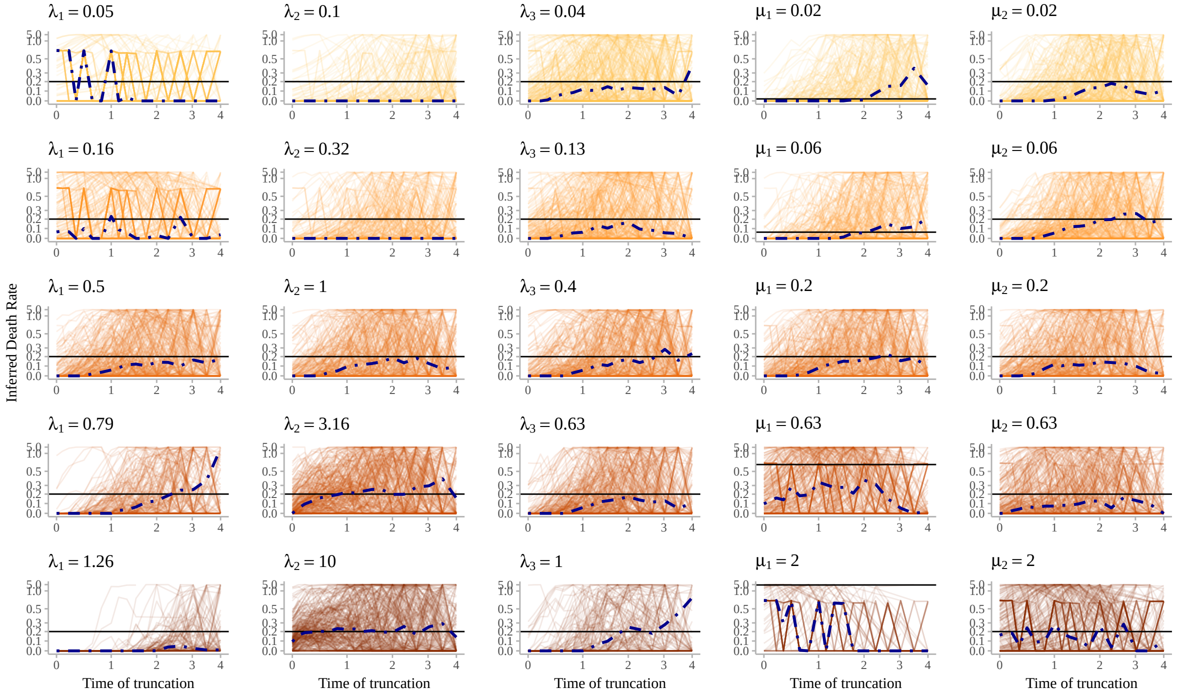


**(b)** Inferred death rates ($\mu$) across truncation lengths and replicates.

**Figure S7. Correspondence between the equivalent time-constant birth-death (BD) rates and inferred birth-death rates on truncated phylogenies generated under the protracted birth-death (PBD) model.** Each column shows the effect of independently modifying a given PBD parameter, with the default value on the third row. For each replicate phylogeny, a classical BD model was fitted on the reconstructed phylogeny and a modified BD model was fitted on the same phylogeny with truncated terminal branches, for increasing truncation lengths. The dot-dashed line corresponds to the median rate across replicates. The horizontal line corresponds to the equivalent time-constant birth or death rate as defined in equation 7.

Note that the default BD model almost always underestimates the equivalent (asymptotic) rates, except for the highest completion rate ($\lambda_2 = 10$), while they are usually recovered with intermediate truncation times.

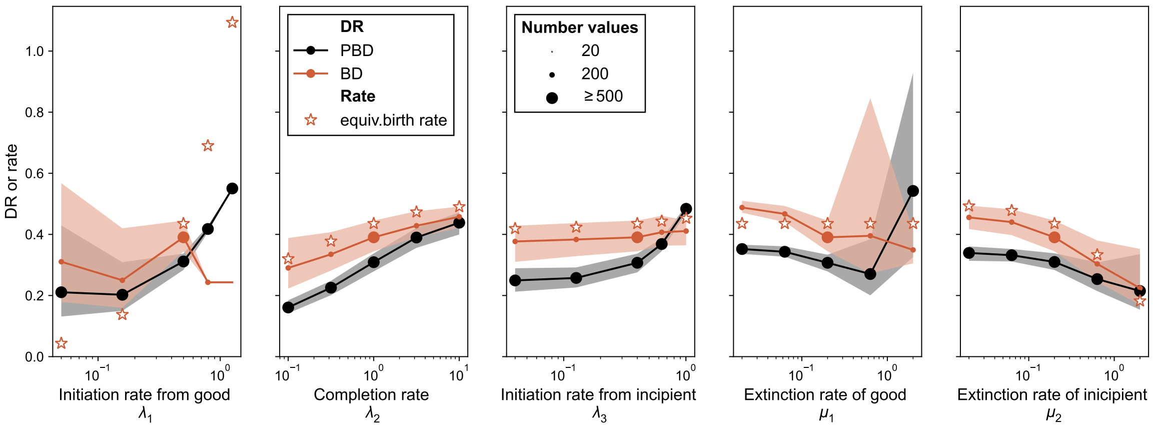


**Figure S8. Diversification rate (DR) statistics on simulated trees under the protracted birth-death (PBD) and birth-death (BD) process with the equivalent rates.** The trees are the same as in figure 5 of the main document. In each column, only one of the PBD parameters varies, with the others held constant (default $\lambda_1 = 0.5, \lambda_2 = 1.0, \lambda_3 = 0.4, \mu_1 = 0.2, \mu_2 = 0.2$). For each set of parameters of the PBD model, trees were also generated under equivalent birth and death rates computed using equation 7 of the main document. On each tree, we computed the median DR across all tips of the tree. The dots and shaded area indicate the median and interquartile ranges of the median DR. The stars indicate the equivalent constant birth rate $\tilde{\lambda}$ as defined in equation 7 of the main document, which is used to generate the BD trees. The DR statistics is defined in Jetz et al. 2012.

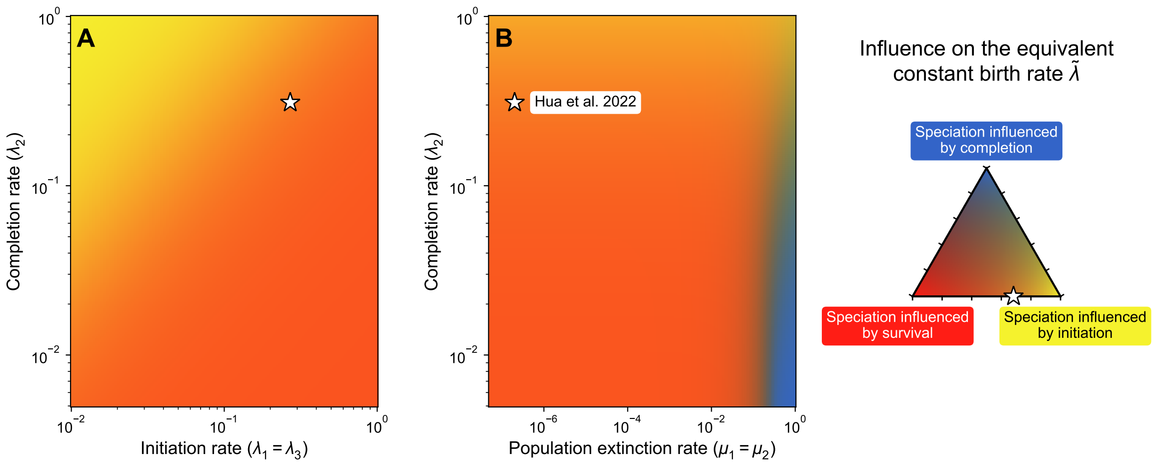


**Figure S9. Relative influence of the parameters of the PBD model on the equivalent constant birth rate, with the values fitted on the phylogeny of Australian Rainbow Skinks in Hua et al. 2022.** Colors indicate which of the PBD process among initiation, completion and population extinction limits the equivalent constant birth rate $\tilde{\lambda}$ most, as a function of (**A**) the initiation and completion rates and (**B**) the population extinction and completion rates, with the color code explained on the triangle in the right. The stars indicate the parameters used as default, taken from the study by Hua et al. 2022 ($b = 0.27, \lambda_2 = 0.31, e = 2e-7$) where the author fitted a ProSSE model (a model slightly different from PBD) to a Rainbow Skinks phylogeny of 41 species. In all cases, the rates of initiation and completion have a positive influence on the birth rate and the rate of population extinction has a negative influence. For the combination of parameters indicated by the stars, we find that the relative influences are 0.682 for $b$, $1.10 \times 10^{-7}$ for $\lambda_2$ and 0.318 for $e$.
